# RORα enforces stability of the T-helper-17 cell effector program

**DOI:** 10.1101/2020.12.15.422921

**Authors:** June-Yong Lee, Jason A. Hall, Maria Pokrovskii, Lina Kroehling, Lin Wu, Dan R. Littman

## Abstract

T helper 17 (Th17) cells regulate mucosal barrier defenses, but also promote multiple autoinflammatory diseases. Although many molecular determinants of Th17 cell differentiation have been described, the transcriptional programs that sustain Th17 cells *in vivo* remain obscure. The transcription factor RORγt is critical for Th17 cell differentiation, but a distinct role of the closely-related RORα, which is co-expressed in Th17 cells, is not known. Here we demonstrate that, although dispensable for Th17 cell differentiation, RORα governs optimal Th17 responses in peripheral tissues. Thus, the absence of RORα in T cells led to significant reductions in both RORγt expression and effector function amongst Th17 cells, due to need for cooperative RORα and RORγt binding to a newly-identified *Rorc* enhancer element that is essential for Th17 lineage maintenance *in vivo*. Altogether, these data point to a non-redundant role of RORα in Th17 lineage maintenance via reinforcement of the RORγt transcriptional program.

## Introduction

T-helper-17 (Th17) cells and related IL-17-producing (Type-17) lymphocytes are abundant at epithelial barrier sites (Honda and Littman, 2016). Their signature cytokines, IL-17A, IL-17F and IL-22, mediate an antimicrobial immune response and also contribute to wound healing and regeneration of injured tissues upon bacterial and fungal infection (Brockmann et al., 2017; Honda and Littman, 2016; Song et al., 2015). However, these cells are also key drivers of multiple chronic inflammatory diseases, including autoimmune diseases and inflammatory bowel disease (IBD), and they have also been implicated in carcinogenesis (Patel and Kuchroo, 2015; Stockinger and Omenetti, 2017). Ultimately, a better understanding of Type-17 regulatory mechanisms may uncover effective therapeutic strategies aimed at treating chronic inflammatory diseases and reducing cancer incidence.

The differentiation of Th17 cells and their ability to produce signature cytokines depend upon induction of the nuclear receptor (NR) transcription factor RAR-Related Orphan Receptor-gamma t (RORγt). RORγt is required for the differentiation of both homeostatic Th17 cells, such as those that regulate commensal microbiota at mucosal barriers, and pro-inflammatory Th17 cells, whose dysregulation results in autoimmune and chronic inflammatory diseases. Therefore, identification of the context-dependent requirements for RORγt expression may facilitate understanding and therapeutic control of inflammatory immune responses. Studies conducted by our group and others have identified some of the *trans*-acting factors necessary for regulating transcription of *Rorc(t)* in Th17 cells (Ciofani et al., 2012; Durant et al., 2010; Schraml et al., 2009). However, the genomic *cis*-regulatory elements that control expression of RORγt in Th17 cells in vivo have been only partially characterized (Chang et al., 2020; Tanaka et al., 2014).

RORγt was initially characterized as the “master regulator” of the Th17 effector program.However, another ROR family transcription factor, RORα, is also upregulated during Th17 cell differentiation, can direct expression of IL-17 (Huh et al., 2011), and was reported to contribute to Th17 cell function (Castro et al., 2017; Yang et al., 2008). Our transcriptional regulatory network analysis of Th17 cells also identified RORα as a key Th17-promoting transcription factor (TF) (Ciofani et al., 2012; Miraldi et al., 2019). By exploring the divergent effects of RORα and RORγt in Th17-driven autoimmune pathogenesis, we found that RORα is crucial for the functional maintenance of the Th17 program, despite exerting a relatively minor influence during differentiation of these cells. Thus, there was reduced accumulation of Th17 cells devoid of RORα in inflamed tissues, which manifested as a dampened pathogenic program. Analysis of chromatin occupancy and accessibility revealed that RORα binds to an enhancer element within the *Rorc* (gene for RORγ and RORγt) locus and positively regulates RORγt expression during chronic autoimmune inflammation. Taken together, these findings suggest that RORα functions as a key regulator for the Th17 effector program through direct regulation of sustained RORγt expression during chronic inflammation.

## Results

### RORα and RORγt are differentially required for Th17-mediated EAE pathogenesis

Although it is established that RORγt is required for Th17 cell differentiation, it has been reported that RORα can partially compensate for RORγt deficiency to promote Th17-dependent experimental autoimmune encephalomyelitis (EAE) (Yang et al., 2008). To study whether these nuclear receptors exert distinct functions in Th17 cells, we studied mice harboring conditional deletions of *Rorc* and/or *Rora* in T cells. In line with previous studies, EAE disease was undetectable (10/18) or mild (8/18) in *CD4^Cre^Rorc^fl/fl^* (T_GKO_) mice, compared to littermate *CD4^Cre^Rorc wt* (T_WT_) animals, which uniformly developed disease following immunization with myelin oligodendrocyte glycoprotein (MOG) in complete Freund’s adjuvant (CFA) and pertussis toxin (Ptx) injection (Figures 1A-C). To determine whether T_GKO_ cells were able to differentiate into Th17 cells in a setting permissive to fulminant EAE disease, we induced EAE in lethally-irradiated Rag1 deficient mice that had been reconstituted with an equal number of isotype-marked CD45.1/2 T_WT_ and CD45.2 T_GKO_ bone marrow cells. In this context, although all mice developed severe EAE, only cells of wild type origin were found to produce IL-17A in the draining lymph nodes (DLN) and spinal cord (SC). Conversely, the proportions of IFNγ-producing cells were similar among T_WT_ and T_GKO_ CD4^+^CD44^+^ T cells in DLN and SC, demonstrating that T_GKO_ cells retained the capacity to acquire effector functions (Figure S1A-B).

**Figure 1.**
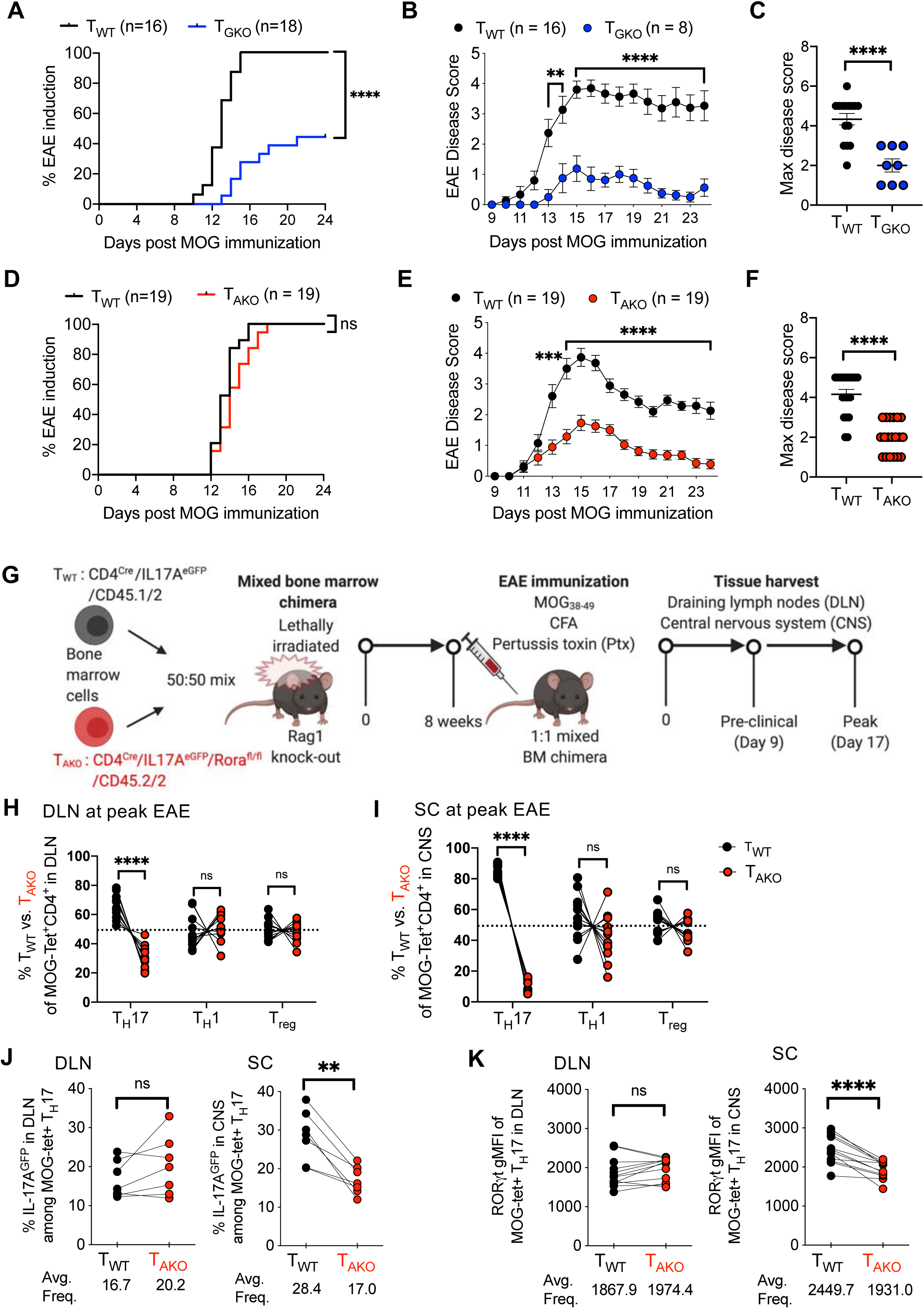
Divergent roles of RORγt and RORα in the differentiation and maintenance of pathogenic Th17 cells in autoimmune encephalomyelitis (EAE). **(A-C)** EAE frequency and severity in T cell-specific RORγt knock-out (T_GKO;_ *CD4^Cre^Rorc^fl/fl^*; n=18) and WT (*CD4^Cre^*; n=16) mice. Time course of EAE incidence **(A)** and mean daily disease score of symptomatic mice **(B)**; maximum disease score of EAE symptomatic mice **(C)**. Summary of 3 experiments. **(D-F)** EAE frequency and severity in T cell-specific RORα knock-out (T_AKO_; *CD4^Cre^Rora^fl/fl^*; n=19) and WT (*CD4^Cre^*; n=19), as in (A-C). Time course of EAE incidence **(D)** and mean daily disease score of symptomatic mice **(E)**; maximum disease score of EAE symptomatic mice **(F)**. Summary of 3 experiments. **(G)** Schematic of EAE induction in CD45.1/2 T_WT_ and CD45.2/2 T_AKO_ 50:50 (T_WT_/T_AKO_) mixed bone marrow (BM) chimeras. **(H and I)** Percent of T_WT_ and T_AKO_ cells of the indicated T cell phenotypes among MOG-tetramer^+^CD4^+^ T cells from draining lymph node (DLN; **H**) or spinal cord (SC; **I**) of T_WT_/T_AKO_ BM chimera at peak of EAE. Each phenotypic program was determined by the specific transcription factor expression by FACS (Th17: RORγt^+^FoxP3^Neg^CD44^hi^CD4^+^ T cells, Th1: T-Bet^+^RORγt^Neg^ FoxP3^Neg^ CD44^hi^ CD4^+^ T cells, Treg: FoxP3^+^CD44^hi^CD4^+^ T cells). **(J)** Percent of IL-17A^eGFP+^ cells among MOG-tetramer^+^CD4^+^RORγt^+^ Th17 cells from DLN (left) or SC (right) of T_WT_/T_AKO_ BM chimera at peak of EAE. **(K)** RORγt gMFI (geometric mean fluorescence intensity) level of MOG-tetramer^+^CD4^+^RORγt^+^ Th17 cells from DLN (left) or SC (right) of T_WT_/T_AKO_ BM chimera at peak of EAE. (A and D) Statistics were calculated by log-rank test using the Mantal-Cox method. (B and E) Statistics were calculated using the two-stage step-up method of Benjamini, Krieger and Yekutieliun. Error bars denote the mean ± s.e.m. (C and F) Statistics were calculated using the unpaired sample T test. Error bars denote the mean ± s.e.m. (E-I) Statistics were calculated using the paired sample T test. ns = not significant, *p < 0.05, **p < 0.01, ***p < 0.001, ****p < 0.0001. (H-K) Data combined from three experiments with 12 BM chimera mice. See also Figure S1.

In contrast to T_GKO_ mice, mice with T cell-specific ablation of *Rora* (*CD4^Cre^Rora^fl/fl^* (T_AKO_)) readily developed EAE (Figure 1D); however, disease severity was substantially milder than in control, littermate *CD4^Cre^Rora wt* (T_WT_) animals (Figures 1E and 1F). To probe the intrinsic role of RORα in pathogenic Th17 cell differentiation, we employed a 1:1 mixed bone marrow chimera strategy similar to that described above (Figures 1G and S1C). Notably, each donor strain also harbored an *Il17a^eGFP^* reporter allele, in order to facilitate examination of myelin-specific Th17 cells using MOG-specific MHC class II (I-A^b^-MOG_38-49_) tetramers (MOG-tet) (Figures 1G and S1D). Assessment in the DLN at the peak of EAE revealed a modest role for RORα in the differentiation of pathogenic Th17 cells, with an almost 2-fold reduction in the frequency of CD45.2/2 T_AKO_ effector Th17 (Foxp3^neg^RORγt^+^CD4^+^) cells relative to CD45.1/2 T_WT_ counterparts (Figures 1H and S1D). By contrast, the proportions of T-effector cells that exclusively expressed the Th1 lineage transcription factor, T-bet, or the regulatory T cell (Treg) lineage transcription factor, FoxP3, were roughly equivalent between the T_AKO_ and T_WT_ populations (Figures 1H and S1D). Strikingly, further skewing (8.2-fold reduction) of the T_AKO_ population relative to wild-type cells was observed among RORγt^+^ Th17 cells in the SC (Figures 1I and S1E). Nevertheless, incorporation of the nucleoside analog EdU indicated that differentiating RORγt^+^ Th17 T_AKO_ effector cells proliferated similarly to their T_WT_ counterparts during the preclinical stage of disease (Figure S1F). Moreover, expression of the S-phase nuclear antigen, Ki67, remained similar in T_AKO_ and T_WT_-Th17 cells located in both the DLNs and SC throughout clinical stages of disease, suggesting that RORα does not regulate accumulation of Th17 cells in the SC via proliferation (Figures S1G and S1H). In concert with their lack of accumulation, MOG-tet^+^ Th17 T_AKO_ cells also exhibited signs of functional impairment in the SC, but not in the DLN, including reduction in proportion of cells expressing the *Il17a^GFP^* reporter and consistent decrease in the mean fluorescence intensity of RORγt expression (Figures 1J and 1K). These data suggest that while RORα is unable to mediate strong Th17 pathogenicity in the absence of RORγt expression, it maintains a prominent role in the regulation of the Th17 effector program.

### RORα is required for a sustained mucosal Th17 response

To address whether the role of RORα in Th17 responses can be generalized, we orally vaccinated co-housed littermate T_WT_ and T_AKO_ mice with an attenuated double mutant (R192G/L211A) form of the heat-labile enterotoxin (dmLT) of enterotoxigenic *Escherichia coli*, which induces a robust antigen-specific mucosal Th17 response (Fonseca et al., 2015; Hall et al., 2008) (Figure 2A). Following two rounds of vaccination, dmLT-specific (I-A^b^-dmLT_166-174_ tetramer positive) cells were readily detectable in the small intestinal lamina propria (SILP) of T_WT_ and T_AKO_ mice (Figure S2A). Yet, both the proportion and number of the dmLT-specific Th17 cells were significantly reduced in T_AKO_ mice (Figures 2B, 2C and S2B). Although this reduction was accompanied by a significant concomitant increase in the frequency of dmLT-specific Th1 cells within the SILP of T_AKO_ mice, both mutant and wildtype counterparts harbored similar numbers of dmLT-Th1 cells, suggesting that only the Th17 component of the effector T-cell response was impaired (Figures 2B, 2C and S2B). Amongst the dmLT-specific Th17 cells, the level of RORγt expression, as well as the frequency of RORγt^+^ cells that expressed CCR6, a RORγt-dependent chemokine receptor, were also significantly reduced in T_AKO_ cells, reinforcing the notion that both RORα and RORγt are required to program and maintain optimal Th17 function (Figures 2D, 2E, S2C and S2D).

**Figure 2.**
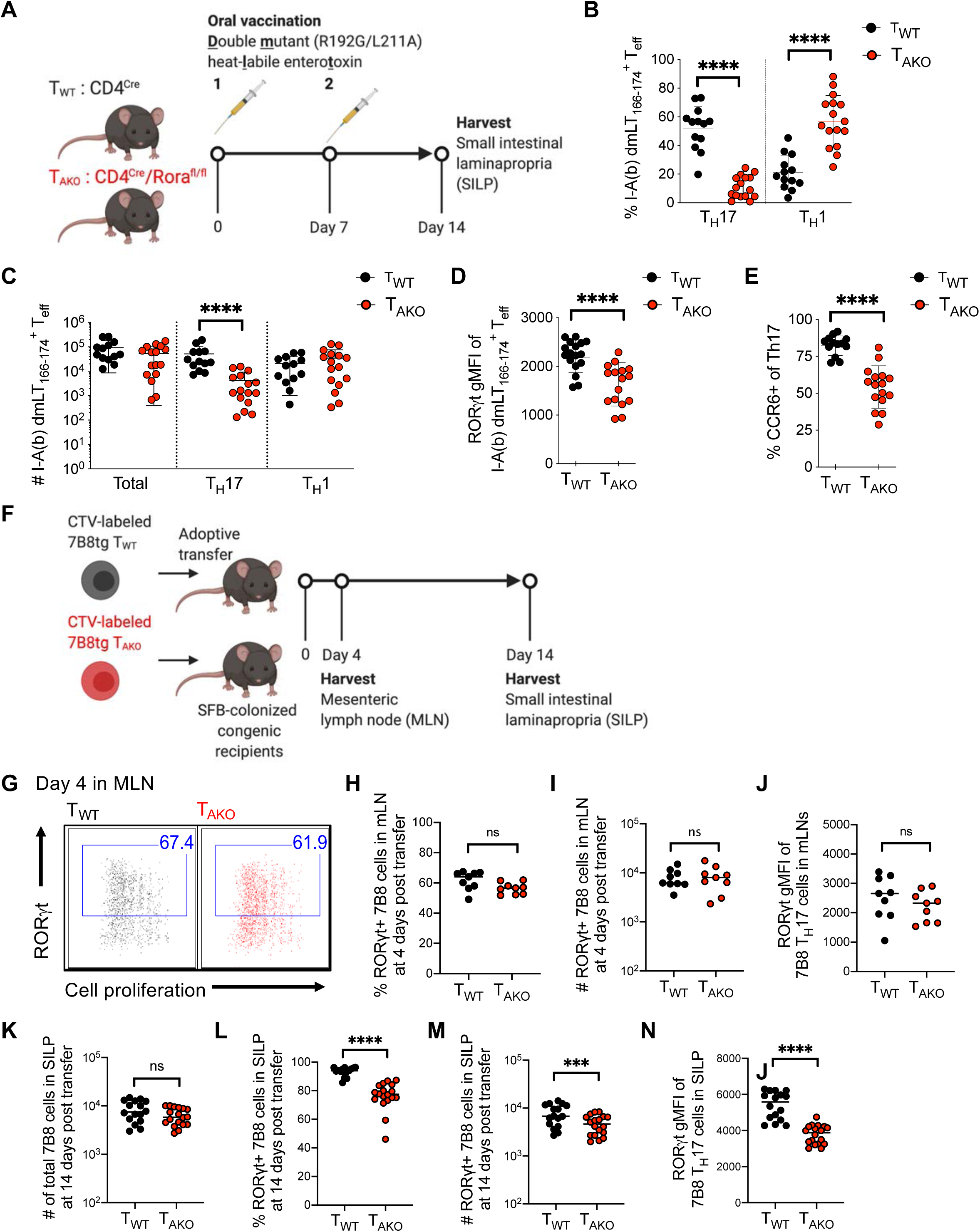
RORα drives sustained mucosal Th17 cell responses. **(A-E)** Oral vaccination of littermate T_WT_ and T_AKO_ mice with an attenuated double mutant (LT R192G/L211A) of the heat-labile enterotoxin of enterotoxigenic *Escherichia coli*, previously shown to induce a robust Th17 response. **(A)** Experimental scheme to examine the role of *Rora* in mucosal Th17 responses. **(B and C)** The proportion **(B)** and absolute number **(C)** of dmLT-specific Th17 and Th1 cells. Phenotypes were determined by FACS profiles for specific transcription factors (Th17: RORγt^+^FoxP3^Neg^CD44^hi^CD4^+^ T cells, Th1: T-Bet^+^RORγt^Neg^ FoxP3^Neg^ CD44^hi^ CD4^+^ T cells, Treg: FoxP3^+^CD44^hi^CD4^+^ T cells). Data combined from three experiments with T_WT_ (n=13) and T_AKO_ (n=16) littermates. **(D)** RORγt gMFI of dmLT-specific Th17 cells. **(E)** Percentage of dmLT-specific Th17 cells expressing CCR6. **(F-N)** RORα deficiency impairs SFB-specific Th17 cell accumulation in SILP. **(F)** Experimental scheme to examine SFB-specific Th17 cell differentiation and effector function of 7B8tg T_WT_ and T_AKO_ in SFB-colonized hosts. **(G-J)** Characterization of donor-derived T_WT_ (n=9) and T_AKO_ (n=9) 7B8tg cells in recipients’ mesenteric lymph nodes (MLN) at 4 days post-adoptive transfer. Flow cytometric analysis of RORγt^+^ Th17 cell differentiation and expansion, monitored by Cell Trace Violet (CTV) dilution **(G)**, and frequency **(H)**, absolute number **(I)** and ROR*γ*t gMFI level **(J)** of RORγt-expressing 7B8tg cells. Data combined from two experiments. **(K-N)** Characterization of donor-derived T_WT_ (n=16) and T_AKO_ (n=18) 7B8tg cells in recipients’ SILPs at 2 weeks post adoptive transfer. Summary of the total numbers **(K)** of SILP-accumulated 7B8tg cells, and frequency **(L)**, absolute number **(M)** and RORγt gMFI level **(N)** of RORγt expressing 7B8tg cells. Data combined from three experiments. Statistics were calculated using the unpaired sample T test. Error bars denote the mean ± s.e.m. ns = not significant, *p < 0.05, ***p < 0.001, ****p < 0.0001. See also Figure S2.

We additionally examined the role of RORα in the differentiation and maintenance of ileal homeostatic Th17 cells induced by segmented filamentous bacteria (SFB). This system allows for study of temporal regulation of Th17 cell differentiation, beginning with priming and proliferation in the draining mesenteric lymph node (MLN) and continuing with expansion and cytokine production in the lamina propria (Sano et al., 2015). T_WT_- or T_AKO_ mice were backcrossed with transgenic mice expressing a TCR (7B8tg) specific for a dominant epitope of SFB (Yang et al., 2014). Naïve 7B8tg T cells from these animals were labeled with Cell Trace Violet (CTV) and adoptively transferred into isotype-distinct hosts colonized with SFB (Figure 2F). Assessment of donor-derived T cells in the intestine-draining MLN revealed that CTV dilution and RORγt induction were similar between T_WT_ and T_AKO_ 7B8tg cells (Figures 2G-J), consistent with the notion that RORα is dispensable for commitment to the Th17 program. Accordingly, similar numbers of T_AKO_ and T_WT_ 7B8tg T cells were recovered two-weeks post-transfer from the terminal ileum section of the SILP, where SFB resides (Figure 2K). However, based on RORγt expression, there was a significant decrease in the proportion and total number of Th17 cells among T_AKO_ compared to T_WT_ 7B8tg T cells (Figures 2L, 2M and S2E), and the RORγt MFI was also reduced in the mutant T cells (Figure 2N and S2F). Altogether, our results indicate that RORα confers the ability of T helper cells to mount a sustained Th17 cell response in target tissues.

### RORα is required for maintenance of the pathogenic Th17 program in the central nervous system

To investigate the molecular mechanism by which RORα regulates the Th17 program, T_WT_ and T_AKO_ Th17 cells were isolated from the DLN and SC of 3 separate cohorts of mixed chimeric mice based on their IL17A^eGFP^ expression (see Figure 1G) at the peak of EAE disease, and their transcriptomes were sequenced (RNA-Seq)(Figures S3A-C). Based on the number of differentially expressed (DE) genes, *Rora* deficiency impacted the Th17 program more profoundly in the SC than in the DLN. At a false discovery rate of 1%, there were 33 DE genes in the DLN, but 845 genes in the SC (Figures 3A, S3B and S3C). The most saliently affected gene in both differentiating (DLN) and effector (SC) T_AKO_-Th17 cells, *Bhlhe40*, was previously found to be required in both Th1 and Th17 cells for manifestation of EAE (Lin et al., 2016). T_AKO_-Th17 cells from the SC also exhibited significant reductions in transcripts encoding proteins that are prominent cell-intrinsic drivers of autoimmune pathogenesis, including *Csf2* (Codarri et al., 2011; El-Behi et al., 2011), *Il1r1* (Shouval et al., 2016), and *Il23r* (Abdollahi et al., 2016; Duerr et al., 2006; Gaffen et al., 2014; Hue et al., 2006) (Figure 3B). Indicative of the sweeping effect that loss of RORα engendered on gene expression at the site of disease, *Rorc*, which encodes RORγt, was markedly reduced in T_AKO_-Th17 cells from the SC, but not from DLN, consistent with reduced expression of direct RORγt target genes (Figure 3C). Thus, combined with the consistent, albeit modest, reduction in protein expression of RORγt in T_AKO_-Th17 cells at effector sites, including the SC and SILP (Figures 1K, 2D and 2N), these findings raise the possibility that RORα reinforces RORγt expression in effector Th17 cells.

**Figure 3.**
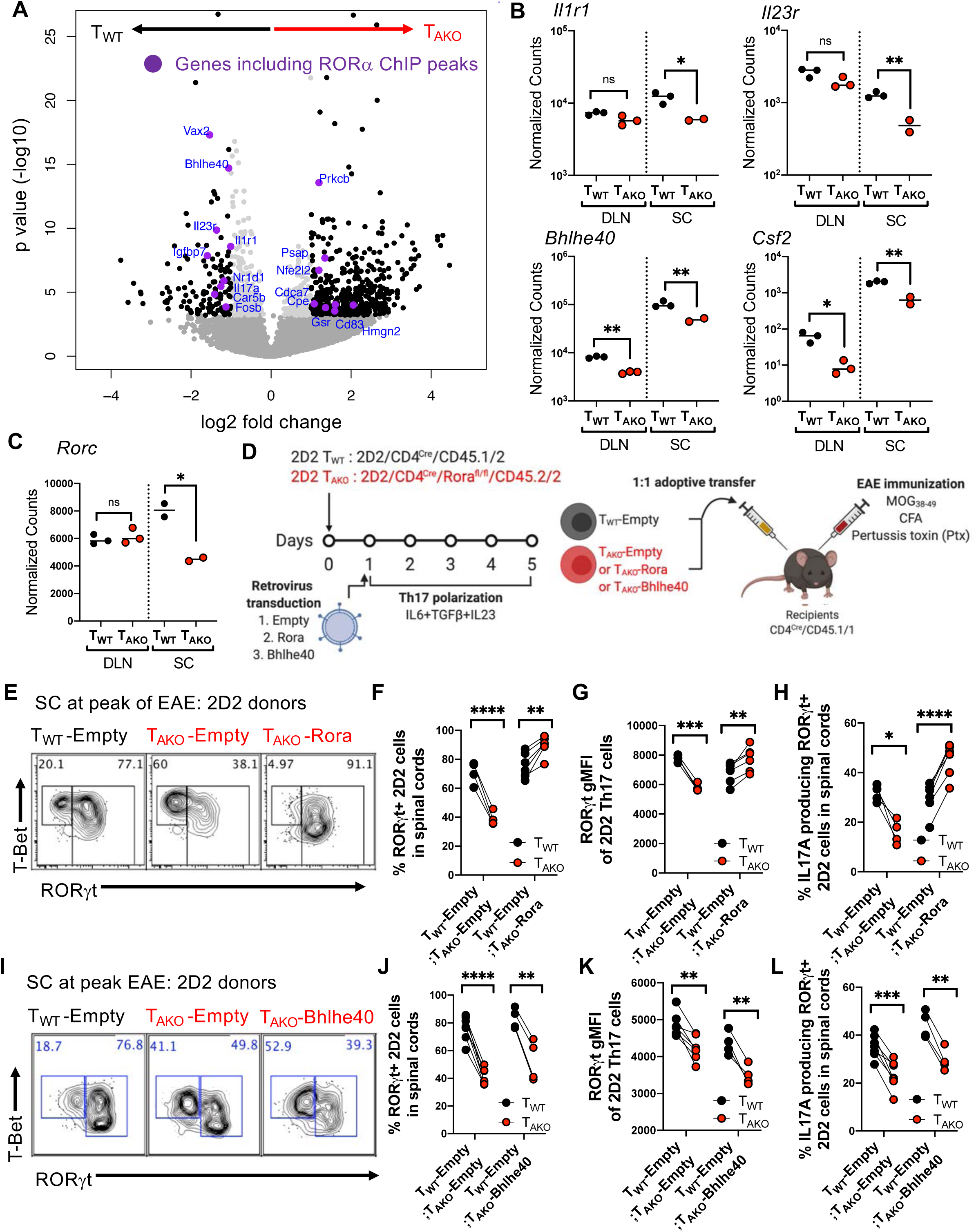
RORα stabilizes the Th17 transcriptional program in effector tissues. **(A-C)** RNA-Seq result of T_WT_ and T_AKO_ Th17 cells, isolated as IL17A^eGFP^-expressing T cells from the DLN and SC of 3 separate cohorts of mixed BM chimera mice at peak of EAE. **(A)** Volcano plot depicting differentially expressed (DE) genes of T_WT_ versus T_AKO_ IL17A^eGFP+^ Th17 cells from the SC. Black dots are significant DE genes. DE genes were calculated in DESeq2 using the Wald test with Benjamini-Hochberg correction to determine the false discovery rate (FDR < 0.01). Purple dots highlight genes that include RORα ChIP-Seq peaks within 10kb of the gene body. **(B and C)** Normalized counts of autoimmune disease-associated (*Il1r1, Il23r, Bhlhe40*), pathogenic (*Csf2*) genes **(B)** and *Rorc* **(C)** in T_WT_ and T_AKO_ *Il17a^eGFP^*^+^ Th17 cells from the DLN (T_WT_ (n = 3) and T_AKO_ (n = 3)) and SC (T_WT_ (n = 3) and T_AKO_ (n = 2)) at peak of EAE. Statistics were calculated using the unpaired sample T test. ns = not significant, *p < 0.05, **p < 0.01. **(D)** Experimental scheme to examine the role of RORα and BHLHE40 in maintenance of the auto-reactive effector Th17 program in inflamed SC during EAE. 2D2tg T_WT_ (CD4^Cre^/CD45.1/2) or T_AKO_ (CD4^Cre^/Rora^fl/fl^/CD45.2/2) cells were retrovirally transduced with *Rora* or *Bhlhe40* or control (Empty) vector, then in vitro polarized to Th17 cells (with IL-6+TGF-β+IL-23) for 5 days. The polarized T_WT_ and T_AKO_ 2D2 cells were combined 1:1 and transferred into recipients (CD4^Cre^/ CD45.1/1) followed by EAE induction (MOG + CFA + Pertussis toxin immunization). **(E)** Flow cytometry analysis of RORγt and T-bet expression of T_WT_, *Rora*-deficient (T_AKO_ -Empty) and *Rora*-reconstituted (T_AKO_-Rora) 2D2 cells in SC at peak of EAE. **(F and G)** Frequency **(F)** and RORγt gMFI **(G)** of RORγt^+^ 2D2tg cells amongst donor T_AKO_-Empty or T_AKO_-Rora 2D2tg cells compared to the T_WT_-Empty in spinal cord at peak of EAE. **(H)** Frequency of indicated IL-17A-producing donor-derived 2D2tg-Th17 cells in SC at peak of EAE following *ex vivo* PMA/Ionomycin re-stimulation. **(I)** Flow cytometry analysis of RORγt and T-bet expression of T_WT_ –Empty and T_AKO_ -Empty or Bhlhe40 ectopic expressing (T_AKO_-Bhlhe40) cells in spinal cord at peak of EAE. **(J and K)** Frequency **(J)** and RORγt gMFI **(K)** of RORγt^+^ T_AKO_-Empty or T_AKO_–Bhlhe40 2D2 T_AKO_ cells compared to T_WT_ -Empty. **(L)** Frequency of indicated IL-17A-producing donor-derived 2D2tg-Th17 cells in SC at peak of EAE following *ex vivo* PMA/Ionomycin re-stimulation. (E-H) Summary of 2 experiments, with T_WT_-Empty:T_AKO_-Empty (n = 4) and T_WT_-Empty:T_AKO_-Rora (n = 6) recipients. Statistics were calculated using the paired sample T test. *p < 0.05, **p < 0.01, ***p < 0.001, ****p < 0.0001. (I-L) Summary of 2 experiments, with T_WT_-Empty:T_AKO_-Empty (n = 7) and T_WT_-Empty:T_AKO_-Bhlhe40 (n = 4) recipients. Statistics were calculated using the paired sample T test. *p < 0.05, **p < 0.01, ***p < 0.001, ****p < 0.0001. See also Figure S3.

To further explore this hypothesis, we developed a retroviral reconstitution system with T cells from MOG peptide-specific (2D2) TCR transgenic mice bred to RORα-deficient or wild-type mice. T_AKO_ 2D2 cells were transduced with *Rora* (yielding T_AKO_-Rora cells) or control (T_AKO_-Empty) vectors and were then cultured under Th17 cell differentiation conditions. They were then transferred with an equal number of similarly prepared isotype-marked T_WT_ 2D2 cells transduced with a control vector (T_WT_-Empty) into recipients that were then immunized to induce EAE (Figures 3D). Critically, the *in vitro* differentiated T_AKO_-Rora, T_AKO_-Empty, and T_WT_-Empty 2D2 cells expressed uniform and equivalent levels of ROR*γ*t prior to adoptive transfer (Figure S3D). Yet, recapitulating the endogenous model, the frequency of RORγt^+^ cells amongst T_AKO_-Empty 2D2 cells in the SC at the peak of disease was markedly reduced relative to that of T_WT_-Empty 2D2tg cells (Figures 3E, 3F, S3E, and S3F). Gating on the RORγt^+^ population also revealed a modest, though significant, decline in protein expression intensity, as well as an impaired capacity to produce IL-17A upon mitogenic restimulation (Figures 3G, 3H and S3G). Each of these deficits was reversed in T_AKO_-Rora 2D2tg cells, corroborating an essential role for RORα in maintenance of the Th17 effector program (Figures 3E-H, S3D-G). The pronounced effect of RORα on *Bhlhe40* expression in differentiating and effector Th17 cells suggested that its influence on Th17 stability may act indirectly through BHLHE40, which is a critical regulator of autoreactive T cell pathogenicity (Lin et al., 2016; Lin et al., 2014). However, ectopic expression of BHLHE40, despite rescuing impaired T_AKO_-2D2 cell accumulation (Figures S3H-J), failed to restore Th17 cell numbers or effector functions among 2D2-T_AKO_ cells (Figures 3I-L and S3K). Thus, regulation of BHLHE40 by RORα is not sufficient to direct effector Th17 cell maintenance, suggesting that RORα regulates other genes that are essential for this differentiation program.

### RORα shares genomic binding sites with RORγt

To ascertain whether RORα directly regulates Th17 lineage maintenance, ChIP-Seq of RORα was performed with *in vitro* differentiated Th17 cells generated from RORα -Twin Strep (RORA-TS) tag knockin-in mice. These animals, which possess a Twin-Strep tag immediately upstream of the stop codon of the *Rora* locus, had normal development and immune cell functions, including frequencies of RORα-dependent type2 innate lymphoid cells (ILC2) (Figures S4A and S4B) and induction of both RORα and RORγt during *in vitro* Th17 cell differentiation on par with WT counterparts (Figures S4C and S4D). Alignment of RORα ChIP peaks with our previously published RORγt ChIP-Seq results for *in vitro* polarized Th17 cells (Ciofani et al., 2012) revealed significant overlaps of genome binding loci between RORα and RORγt, including previously reported genes involved in the “pathogenic” Th17 effector program (e.g., *Il17a/f*, *Il23r* and *Bhlhe40*) (Lee et al., 2012) (Figures 3A, 4A and S4E), and gene ontology analysis of the RORα direct target genes also revealed a significant enrichment in Th17 effector functions and Th17-mediated disease pathogenesis (Figure 4B). Notably, RORα also binds to intronic regions of *Rorc* (Figure 4C). To further address the interdependency of RORα and RORγt in binding to target loci, RORα ChIP-Seq was also conducted on Th17-polarized CD4^+^ T cells isolated from RORA-TS mice in which RORγt activity was abolished (RORA-TS-T_GKO_). Although loss of ROR*γ*t expectedly impeded Th17 cell differentiation (Figure S4F), both *Rora* induction and protein expression were comparable between WT and RORA-TS-T_GKO_ cells cultured under Th17 polarizing conditions (Figures S4G and S4H). Nevertheless, the majority of RORα peaks were ablated upon loss of RORγt (Figures 4A and S4E). In contrast, RORγt binding was not adversely affected in Th17-polarized cells that reciprocally lacked ROR*α* (Figure S4I). These findings are concordant with a limited and largely RORγt-dependent role for RORα during in vitro Th17-cell differentiation.

**Figure 4.**
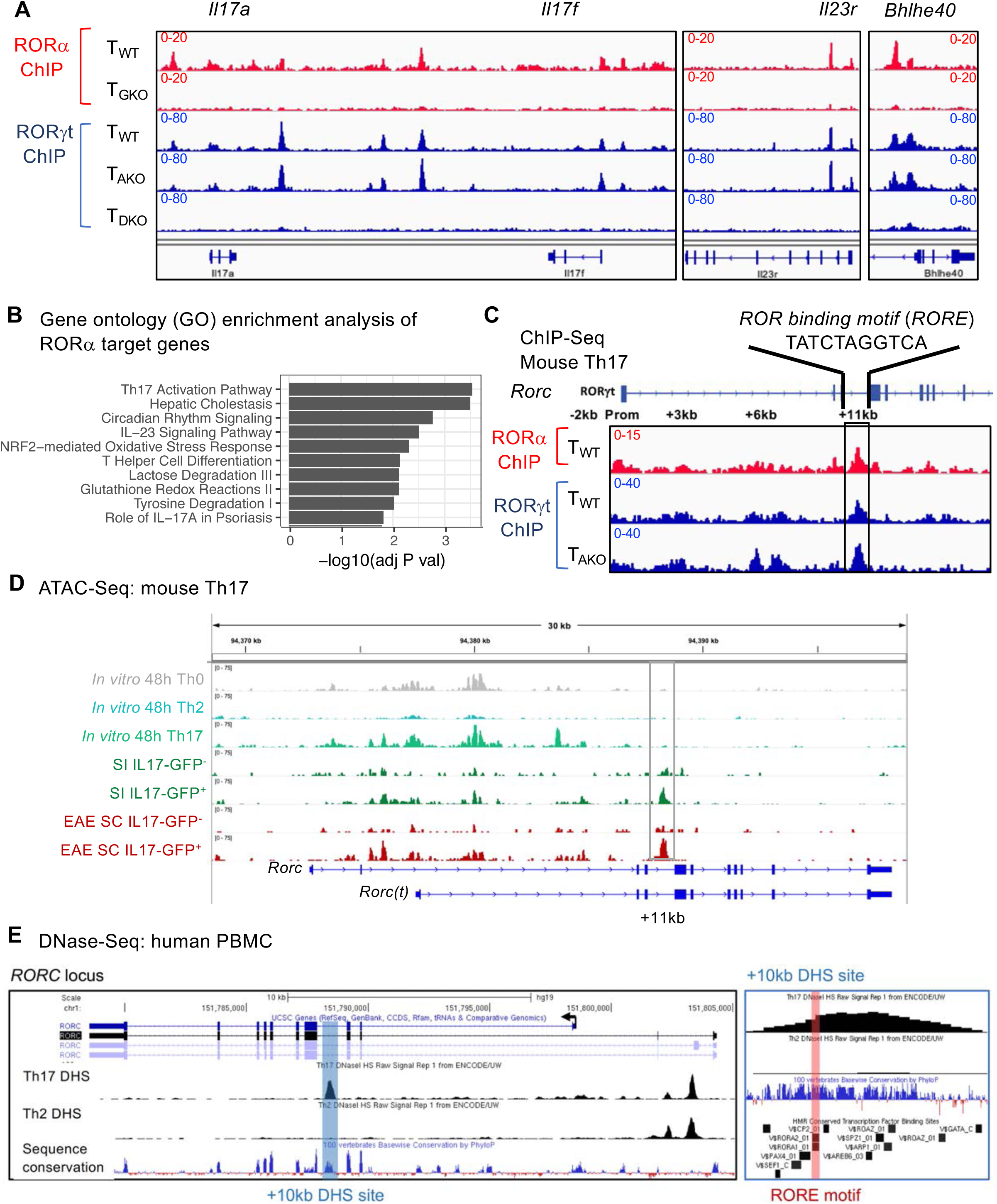
RORα shares genomic binding sites with RORγt in Th17 cells. **(A)** ChIP-Seq tracks of RORγt and RORα within Th17 effector program genes. **(B)** Gene ontology analysis of ROR*α* direct target genes (Peak(s) found within 10kb of gene body). **(C)** ChIP-Seq data exhibiting RORγt and RORα binding to cis-regulatory elements in *Rorc* locus. **(D)** ATAC-Seq data showing open cis-elements in the *Rorc* locus of *in vitro* differentiated or *ex vivo* isolated T cell lineages. Small intestine (SI) or EAE spinal cord (SC) T cells were FACS sorted from *Il17a^eGFP^* mice gated on TCRβ^+^ then either GFP positive or negative. **(E)** UCSC genome browser depicting DNase-Seq on human Th17 (UCSC Accession: wgEncodeEH003020) and Th2 (UCSC Accession: wgEncodeEH000491) from the Encode database aligned with GRCh37/hg19 and the Vertebrate Multiz Alignment & Conservation (100 Species) and HMR Conserved Transcription Factor Binding Sites tracks. *RORC* locus (left) and zoomed +10kb DHS site (right). See also Figure S4.

### The *Rorc(t)* +11kb locus is required for RORα-mediated RORγt expression in tissue-resident Th17 cells

In support of the hypothesis that RORα can directly regulate RORγt expression, ChIP-Seq revealed a significant RORα peak with an embedded ROR response element (RORE) at +11kb from the *Rorc(t)* transcriptional start site in Th17 cells generated *in vitro* (Figure 4C). Alignment with RORγt ChIP-Seq data demonstrated that both family members bind to this region (Figure 4C). Surprisingly, although the assay for transposase-accessible chromatin sequencing (ATAC-Seq) indicated that this region remained closed in *in vitro*-differentiated Th17 cells, it was readily accessible in *ex vivo* IL-17A^+^ Th17 cells sorted from the SILP and both DLNs and SC during EAE (Figure 4D). Moreover, comparison of chromatin accessibility in T-helper lineages enriched from PBMC under the ENCODE Project (Maurano et al., 2012) revealed a prominent syntenic DNase Hypersensitivity Site (DHS) at +10kb from the *RORC* transcription start site (TSS) that was specific to Th17 cells, highlighting that this region constitutes a functionally conserved enhancer in human Type-17 immunity (Figure 4E). Altogether, these data suggest that synergy of RORα and RORγt binding to the intronic RORE following early RORγt induction governs subsequent RORγt stability in Th17 cells *in vivo*.

To functionally interrogate the role of the *Rorc(t)* +11kb cis-element in vivo, we generated transgenic mice with a *Rorc*-containing BAC engineered to have a mCherry reporter at the RORγt translational start site with or without deletion of the +11kb *cis*-element (WT Tg (Rorc(t)-mCherry and Δ+11kb Rorc(t)-mCherry) (Figure 5A). To serve as an internal control, the transgenic mice were bred to *Rorc(t)^GFP^* mice containing a GFP reporter knocked into the endogenous *Rorc(t)* locus (Figure 5A and S5A). Thymocyte development was normal in both WT Tg and Δ+11kb Tg lines, with mCherry expression highest in double positive and early post-selection single positive thymocytes, consistent with known expression patterns of RORγt (He et al., 2000; Sun et al., 2000) (Figure S5B). Within the SILP, a strong correlation between GFP and mCherry expression was also observed in both innate and adaptive Type-17 lymphocytes, which included not only Th17 cells, but also γδT cells and type 3 innate lymphoid cells (ILC3) of WT Tg mice (Figure 5B, 5C and S5C-E). In stark contrast, mCherry activity within the SILP of Δ+11kb Tg mice was lost in each of these populations, suggesting that the +11kb *cis*-element is a bona fide enhancer for all Type-17 lymphocyte lineages in vivo (Figure 5B, 5C and S5C-E). Nevertheless, CD4^+^ T cells isolated from Δ+11kb Tg mice readily expressed mCherry upon *in vitro* Th17 polarization (Figure 5D and 5E). This finding, together with the chromatin accessibility data for *in vitro* polarized Th17 cells (Figure 4D), as illustrated by failure to open chromatin at the +11kb locus, suggests that the +11kb *cis*-element is an essential enhancer for the Type-17 lymphocytes in vivo but is dispensable for thymocyte development and *in vitro* Th17 cell differentiation.

**Figure 5.**
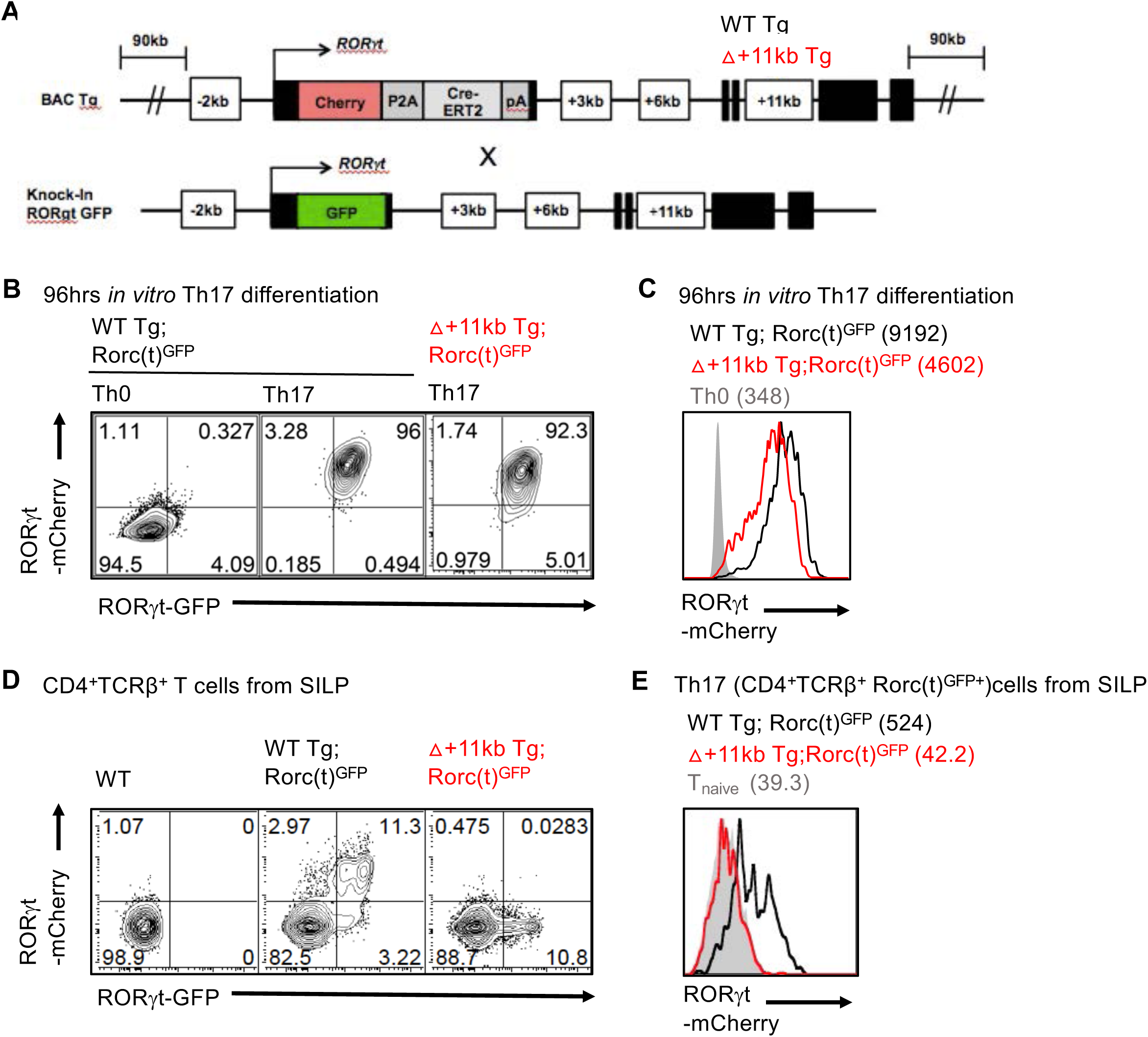
The *Rorc* +11kb cis-element is required for RORγt expression in Th17 cells *in vivo*, but is dispensable for *in vitro* differentiation. **(A**) Schematic depicting endogenous and BAC transgene allele in WT Tg (Rorc(t)-mCherry);Rorc(t)^GFP^ control or +11kb enhancer mutant (Δ+11kb) Tg (Δ+11kb Rorc(t)-mCherry);Rorc(t)^GFP^ mice. **(B and C)** Flow cytometry plots (B) and stacked histogram (C) illustrate Rorc(t)-mCherry reporter expression in *in vitro* differentiated Th17 cells from WT Tg; Rorc(t)^GFP^ or Δ+11kbTg;Rorc(t)^GFP^ mice. Geometric mean fluorescence intensities (gMFI) are included in parentheses. Representative data of three experiments. **(D and E)** Flow cytometry plots (D and stacked histogram (E) illustrates Rorc(t)-mCherry reporter expression in *ex vivo* isolated Th17 (TCRβ^+^RORγt^GFP+^) cells from SILP of WT Tg (Rorc(t)-mCherry);Rorc(t)^GFP^ control or +11kb enhancer mutant (Δ+11kb) Tg (Δ+11kb Rorc(t)-mCherry);Rorc(t)^GFP^ mice. gMFIs are included in parentheses. Representative data of three experiments. See also Figure S5.

To further investigate whether the +11kb conserved noncoding sequence functions via the binding of ROR family TFs in EAE, an optimized Cas9/gRNA RNP transfection approach was utilized to mutate the RORE and preclude RORα and ROR*γ*t binding to the +11kb-enhancer in *in vitro*-differentiated Th17 cells (Figure 6A). Targeting the locus in activated naïve 2D2tg-T cells resulted in nearly 100% editing efficiency, with both indels and deletions that did not exceed 100bps (Figures S5A and S5B). Following *in vitro* Th17 cell polarization with IL-6, TGF-β and IL-23, control gene (sgCtrl) and +11kb-enhancer-targeted (+11kb^ΔRORE^) 2D2tg-Th17 cells were adoptively transferred into wild-type recipients, which were then immunized with MOG peptide to trigger EAE (Figure 6A). Consistent with the inaccessibility the *Rorc* +11kb site in *in vitro* polarized Th17 cells, neither the induction of RORγt, nor the capacity to secrete IL-17A, were affected in the +11kb^ΔRORE^ 2D2tg-Th17 cells at the time of transfer (Figures 6B and 6C). However, by the peak of disease, in comparison to control-targeted counterparts, both the percentage and absolute number of RORγt^+^ +11kb^ΔRORE^ 2D2tg-Th17 cells recovered from the SC sharply declined (Figures 6D-F). Among the residual Th17 cells, RORγt expression levels were also significantly reduced (Figure 6G). These findings reflect the compromised stability of RORγt in T_AKO_ 2D2tg-Th17 cells during EAE (Figures 3E-H). Accordingly, ectopic overexpression of *Rora* restored RORγt expression in T_AKO_ 2D2tg-Th17 cells, but not in T_AKO_ +11kb^ΔRORE^ 2D2tg-Th17 cells (Figures 6H and 6I), consistent with RORα binding to the +11kb enhancer mediating sustained expression of RORγt. Thus, our findings uncover a novel enhancer required for maintenance of the Th17 cell program in tissues and regulated, at least in part, by RORα.

**Figure 6.**
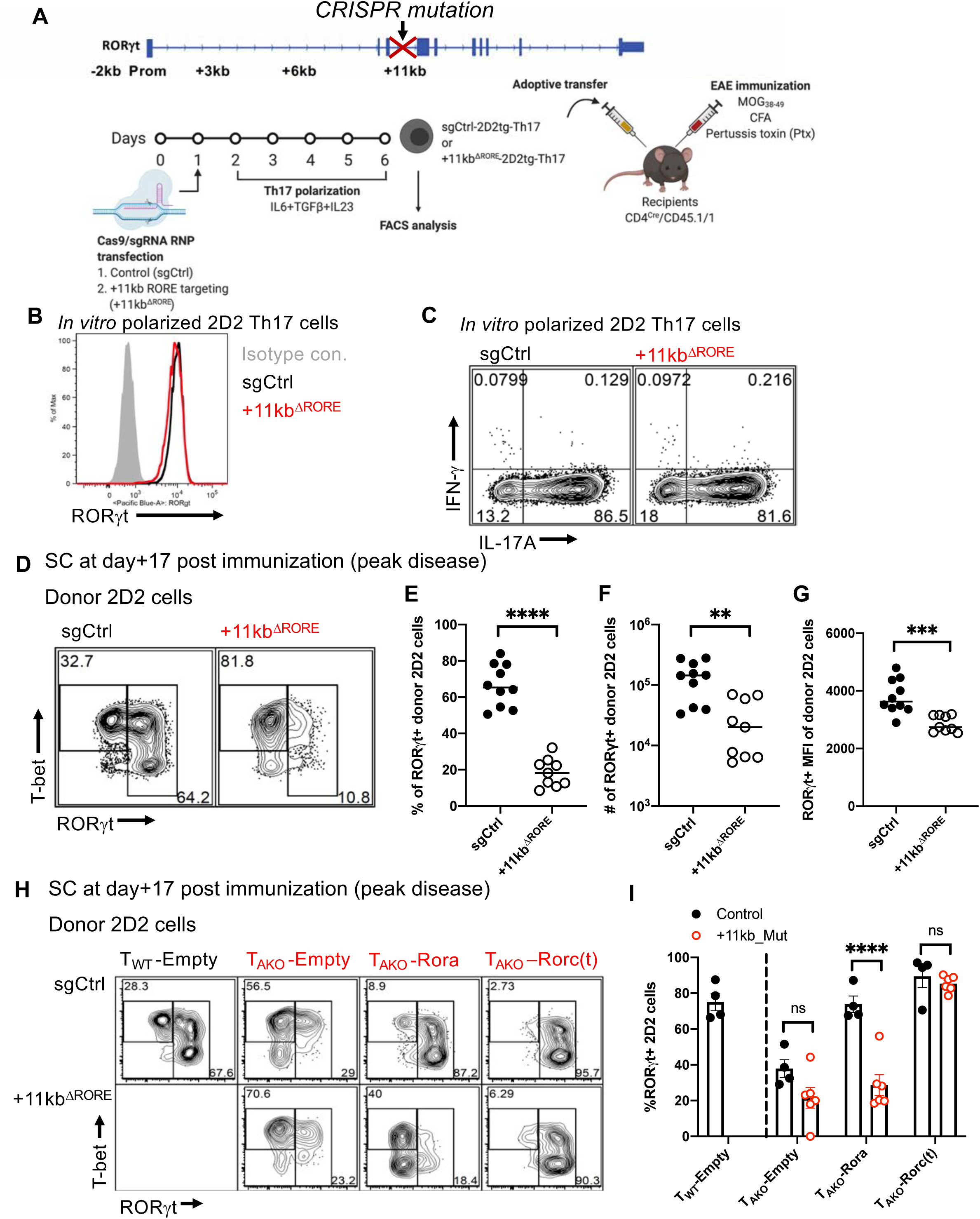
RORα promotes *in vivo* Th17 stability through a conserved enhancer located in the +11kb region of the *Rorc(t)* locus. **(A)** Experimental scheme to interrogate the role of the *Rorc* +11kb element *in vivo.* **(B)** Stacked histogram illustrates RORγt expression in control (sgRNA control; sgCtrl) and *Rorc(t)* +11kb enhancer mutant (sgRNA that target RORE in +11kb cis-element of *Rorc(t)*; +11kb^ΔRORE^) *in vitro* differentiated 2D2tg Th17 cells. **(C)** Representative FACS plots displaying IL-17A and IFNγ production of *in vitro* polarized Th17 sgCtrl or +11kb^ΔRORE^ 2D2tg cells. **(D)** Representative flow cytometry analysis of RORγt and T-bet expression in sgCtrl and +11kb^ΔRORE^ donor-derived 2D2tg cells in SC at peak of EAE. **(E-G)** Frequency (E), number (F) and RORγt gMFI (G) of RORγt-expressing sgCtrl or +11kb^ΔRORE^ 2D2tg cells in SC at peak of EAE. Summary of 2 experiments, with sgCtrl (n = 10) and +11kb^ΔRORE^ (n = 9) recipients. **(H and I)** Flow cytometry analysis of RORγt and T-bet expression (H) and frequency of RORγt expression (I) in sgCtrl or +11kb^ΔRORE^ T_AKO_ donor-derived 2D2tg cells, retrovirally reconstituted with *Rora* or *Rorgt*, in SC at peak of EAE. Summary of 2 experiments, with T_WT_-sgCtrl-Empty:T_AKO_-sgCtrl-Empty (n=4), T_WT_-sgCtrl-Empty:T_AKO_-sgCtrl-Rora (n=4), T_WT_-sgCtrl-Empty:T_AKO_-sgCtrl-Roc(t) (n=4), T_WT_-sgCtrl-Empty:T_AKO_-+11kb^ΔRORE^ -Empty (n=5), T_WT_-sgCtrl-Empty:T_AKO_-+11kb^ΔRORE^-Rora (n=5), T_WT_-sgCtrl-Empty:T_AKO_-+11kb^ΔRORE^ -Roc(t) (n=5) recipients. Statistics were calculated using the unpaired sample T test. Error bars denote the mean ± s.e.m. ns = not significant, *p < 0.05, **p < 0.01, ***p < 0.001, ****p < 0.0001. See also Figure S6.

## Discussion

### Regulation of the Th17 program by RORα and RORγt

Our current study confirms that both RORα and RORγt play important roles in orchestrating Th17 lineage maintenance. Our data suggest that RORα and RORγt may regulate the expression of Th17-associated genes through binding to the same ROREs with their highly similar DNA-binding domains (Cook et al., 2015). This implies that the individual expression levels of RORα and RORγt might be limiting in T cells, leaving ROREs unoccupied, and that expression of both nuclear receptors is required to saturate RORE binding sites and drive maximal ROR-responsive gene expression. Nevertheless, we also observed that expression of RORγt is a prerequisite for RORα binding to the shared RORE. In the absence of RORγt, the T_GKO_ Th17 cells lost most of the genome-wide binding of RORα at the shared target sites. Considering that RORα expression was not impaired upon RORγt deletion, these data are consistent with the model previously proposed by Ciofani et. al, in which RORγt serves as a master switch for Th17 differentiation and creates a feedback pathway that, in turn, stabilizes Th17 commitment (Ciofani et al., 2012). Indeed, we observed that heterozygous loss of RORγt expression impaired optimal in vivo Th17 cell differentiation (Pokrovskii, 2018).

Another possible scenario is that RORα and RORγt may bind to DNA cooperatively. Like all nuclear receptors, ROR proteins have been shown to bind cognate DNA elements as monomers or dimers: as monomers to ROREs containing a single consensus half site (PuGGTCA) immediately preceded by a short A/T-rich region, and as dimers to tandem half sites oriented as palindromes, inverted palindromes, or direct repeats (Giguere, 1999). Indeed, RORα:RORγt heterodimers could possess distinct functional activity compared to monomers or homodimers because of their unique N-terminal trans-activation domains (NTDs) (Giguere, 1999; McBroom et al., 1995). Yet, the precise hierarchical roles of RORα and RORγt in Th17 differentiation and function are unclear and need to be further elucidated.

### Diverse immune functions of RORα

Although our *in vivo* studies indicate a Th17-specific role of RORα during mucosal immune responses to SFB and oral vaccination, as well as in Th17-mediated EAE pathogenesis, RORα has also been found to have important roles in other immune cell types that function in different contexts. RORα is known to be important for development of type 2 innate lymphoid cells (ILC2) (Halim et al., 2012) and for cytokine production by ILC3 (Lo et al., 2019; Lo et al., 2016). In the liver, RORα controls inflammation by promoting macrophage M2 polarization (Han et al., 2017; Kopmels et al., 1992). Among other T cell lineages, it was also shown to have an important function in skin Tregs, where it is required for proper regulation of the immune response in atopic dermatitis (Malhotra et al., 2018). A recent study showed that RORα is involved in mounting Th2 responses during worm infections and allergy (Haim-Vilmovsky,et al., 2019). RORα thus has wide-ranging activities in the regulation of immune responses. Mechanisms by which RORα differentially regulates transcription in these diverse cell types *in vivo* under various immune-contexts remain poorly understood. Furthermore, it is not clear what direct target genes RORα controls in non-Type17 cells that lack RORγt expression. To elucidate the context-dependent, cell type-, and tissue-specific targets of RORα in small numbers of *ex vivo* isolated cells, new and more sensitive technologies will be needed to identify with base-pair precision the regions of DNA that it occupies.

### RORα as a potential drug target for Th17-mediated chronic inflammatory conditions

Chronic inflammation underlies a number of debilitating human diseases including inflammatory bowel disease, multiple sclerosis, psoriasis, and various arthritides (Bamias et al., 2016; Firestein and McInnes, 2017; Netea et al., 2017; Noda et al., 2015). Th17 cells have central roles in many of these diseases. The transcription factor RORγt was initially coined the master regulator of the Th17 program, but targeting RORγt therapeutically is dangerous owing to an enhanced risk of thymoma upon its inhibition (Guntermann et al., 2017; Guo et al., 2016; Liljevald et al., 2016). RORα was also implicated in Th17 functions (Castro et al., 2017; Yang et al., 2008), but its precise role and relationship to RORγt function were not investigated. Exploration of the divergent effects of RORα and RORγt in Th17-elicited autoimmune pathogenesis revealed that RORα is crucial for the functional maintenance of the Th17 program at the site of inflammation despite exerting a relatively minor influence during differentiation in the lymph nodes. During EAE, Th17 cells devoid of RORα are limited in their accumulation in the central nervous system, and those that are present display a dampened pathogenic program. Probing the intersection of ROR binding targets identified by ChIP-Seq with RNA-Seq data obtained from ex vivo isolated RORα-deficient Th17 cells indicated that the majority of RORα targets are shared with ROR*γ*t. Among the most significant were the IL-23 receptor, *Il23r*, and the transcription factor *Bhlhe40*, which are critical for driving Th17 pathogenesis by way of inflammatory T cells having shared Th17 and Th1 features (Harbour et al., 2015; Hirota et al., 2011). Strikingly, RORα was also found bound to a conserved *cis*-regulatory enhancer element in the *Rorc* locus that is crucial for maintenance of RORγt expression in effector Th17 cells *in vivo*. Intriguingly, using our laboratory’s previous transcription factor binding data (Ciofani et al., 2012), Chang et al. recently identified this region (CNS11) in their study of Th17 enhancers, but did not prosecute its function owing to its lack of H3K27Ac marks and weak interaction with p300 (Chang et al., 2020). These data are also consistent with the marginal chromatin accessibility of the +11kb region observed upon *in vitro* differentiation and suggests that a heretofore unidentified factor mediates *in vivo* accessibility of this region. Together, these findings indicate that RORα not only regulates the Th17 program by a means parallel to RORγt, but that it may serve a particularly prominent in vivo function by dynamically reinforcing RORγt expression in the absence of saturating levels of active RORγt.

Natural ligands and synthetic compounds that modulate the function of nuclear receptors have demonstrated tremendous therapeutic potential for multiple clinical conditions (Cheng et al., 2019; Huh et al., 2011; Kojetin and Burris, 2014; Marciano et al., 2014; Moutinho et al., 2019). Our current study, by identifying RORα as a key regulator of the sustained Th17 effector program, suggests that targeting this receptor could be a viable strategy for treating autoimmune pathologies linked to Th17 effector functions in chronically inflamed patient tissues. Furthermore, the involvement of RORα in ILC2 development (Halim et al., 2012; Wong et al., 2012) and Type-2 immune functions (Haim-Vilmovsky,et al., 2019) may provide additional therapeutic opportunities for diseases such as asthma, chronic obstructive pulmonary disease (COPD), and idiopathic pulmonary fibrosis (Gieseck et al., 2018).

However, like other ROR family members, RORα regulates multiple non-immune cell types, in non-inflammatory contexts. For example, *staggerer* mice, which carry a spontaneous deletion in *Rora*, have an underdeveloped cerebellar cortex, with deficiency in granule and Purkinje cells (Gold et al., 2007). RORα has also been linked to neurologic disorders, including autism, in humans (Devanna and Vernes, 2014; Nguyen et al., 2010; Sarachana and Hu, 2013). Significant circadian disruption, described in autistic patients (Hu et al., 2009; Melke et al., 2008; Nicholas et al., 2007), may be related to the role of RORα in regulation of the circadian clock (Jetten, 2009; Kojetin and Burris, 2014). Therefore, a deeper understanding of cell type-specific and context-dependent regulation of RORα is likely needed to inform strategies to combat RORα-associated immune diseases.

In summary, our study has elucidated a non-redundant role of RORα in Th17 lineage maintenance via reinforcement of the RORγt transcriptional program. Further characterization of the interaction of these two nuclear receptors may enable more refined strategies to target specific processes that fuel chronic inflammatory disease.

## Acknowledgements

We thank members of the Littman laboratory for valuable discussions, Sang Y. Kim at the Rodent Genetic Engineering Core (RGEC) of NYU Medical Center (NYULMC) for generation of RORA-TS mice, Adriana Heguy and the Genome Technology Center (GTC) for RNA and ChIP sequencing. We also thank Yasmine Belkaid (NIAID), Oliver Harrison (Benaroya Research Institute) and Timothy Hand (University of Pittsburgh) for providing the specific MHCII (I-A^b^-dmLT_166-174_) tetramer and for helpful discussions surrounding the dmLT vaccine data. Lyophilized dmLT was generously provided by Elizabeth Norton at Tulane University. The GTC and RGEC are partially supported by Cancer Center Support grant P30CA016087 at the Laura and Isaac Perlmutter Cancer Center. This work was supported by an HHMI Fellowship of the Damon Runyon Cancer Research Foundation 2232-15 (J.-Y.L.), a Dale and Betty Frey Fellowship of the Damon Runyon Cancer Research Foundation 2105-12 (J.A.H), the Howard Hughes Medical Institute (D.R.L.), the Helen and Martin Kimmel Center for Biology and Medicine (D.R.L.), and National Institutes of Health grants R01AI121436 and R01DK103358 (D.R.L.).

## Author Contributions

J.-Y.L., J.A.H, and D.R.L. conceived the project; J.-Y.L. and J.A.H. performed the experiments; M.P. investigated cis-regulatory elements of *Rorc(t)* locus; L.K. performed bioinformatic analyses; L.W. contributed to antibody generation and purification; J.-Y.L., J.A.H, M.P. and D.R.L. wrote the manuscript.

## STAR Methods

### KEY RESOURCE TABLE

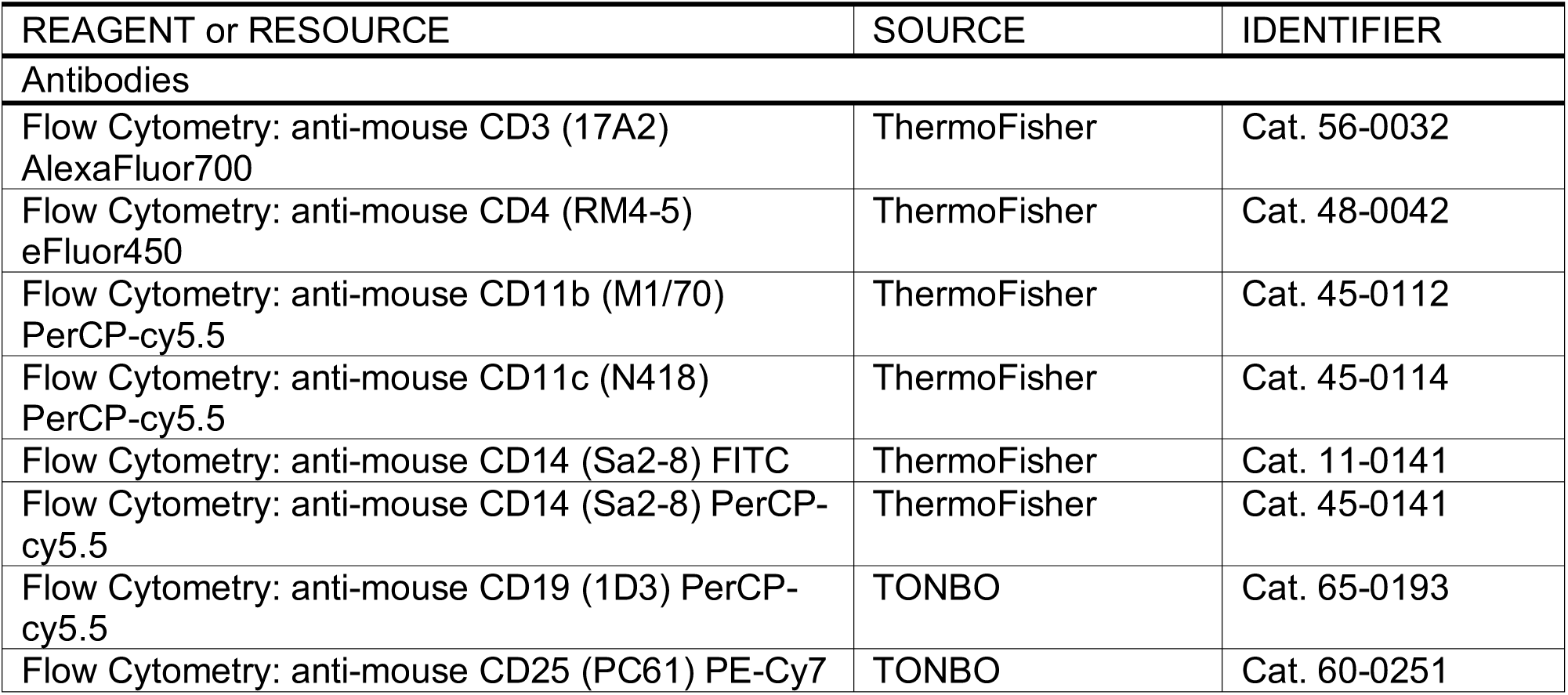
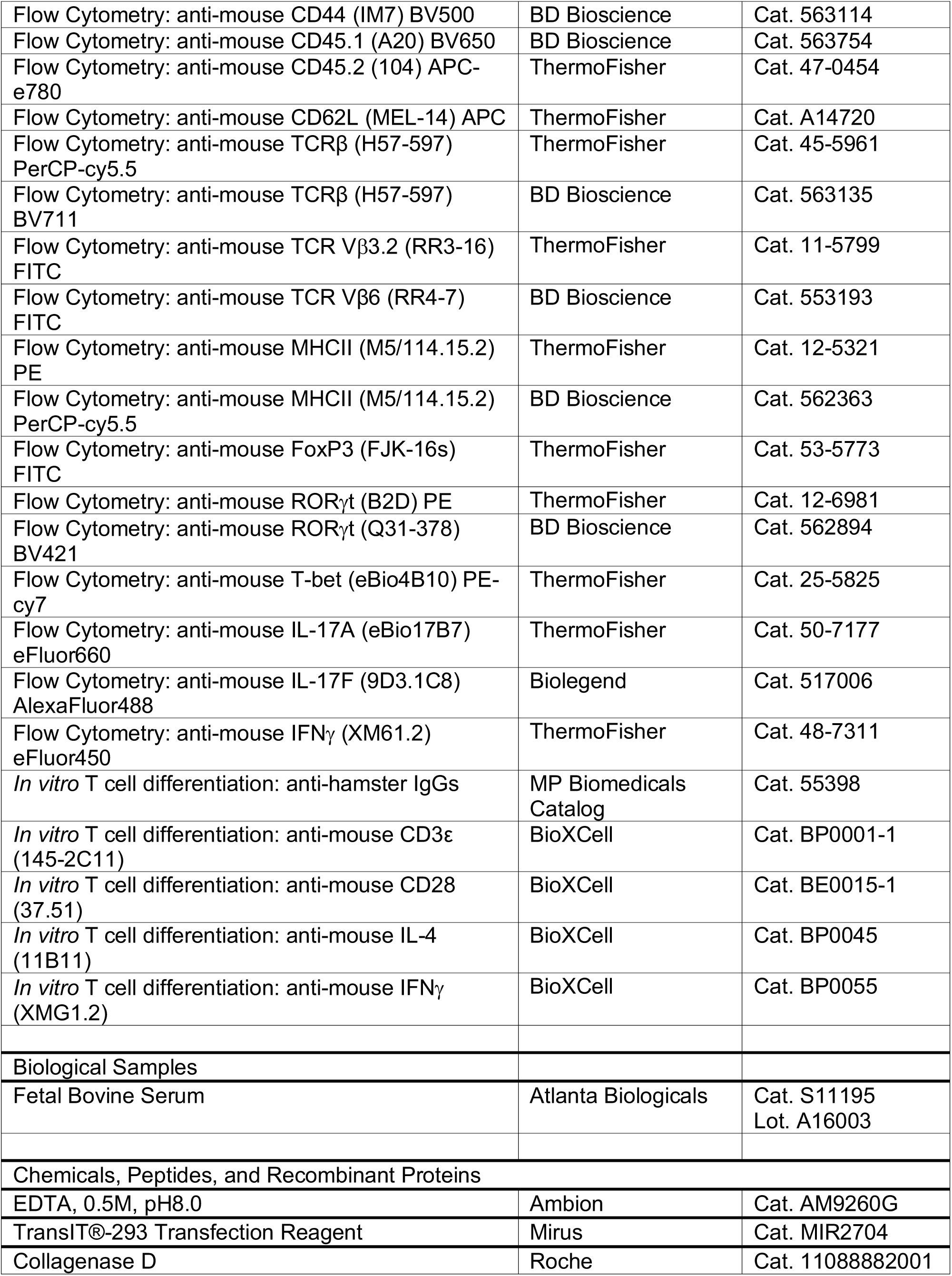
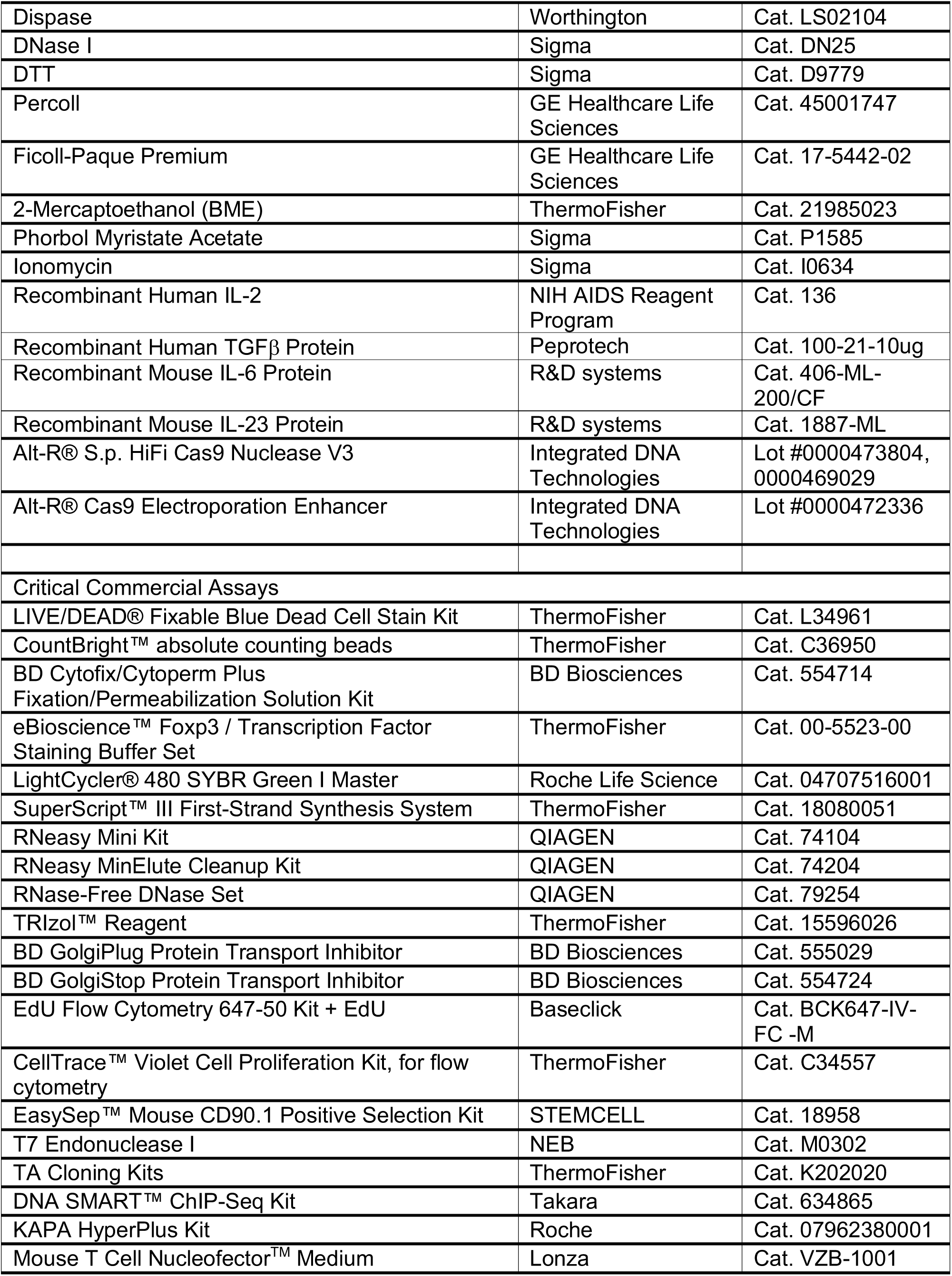
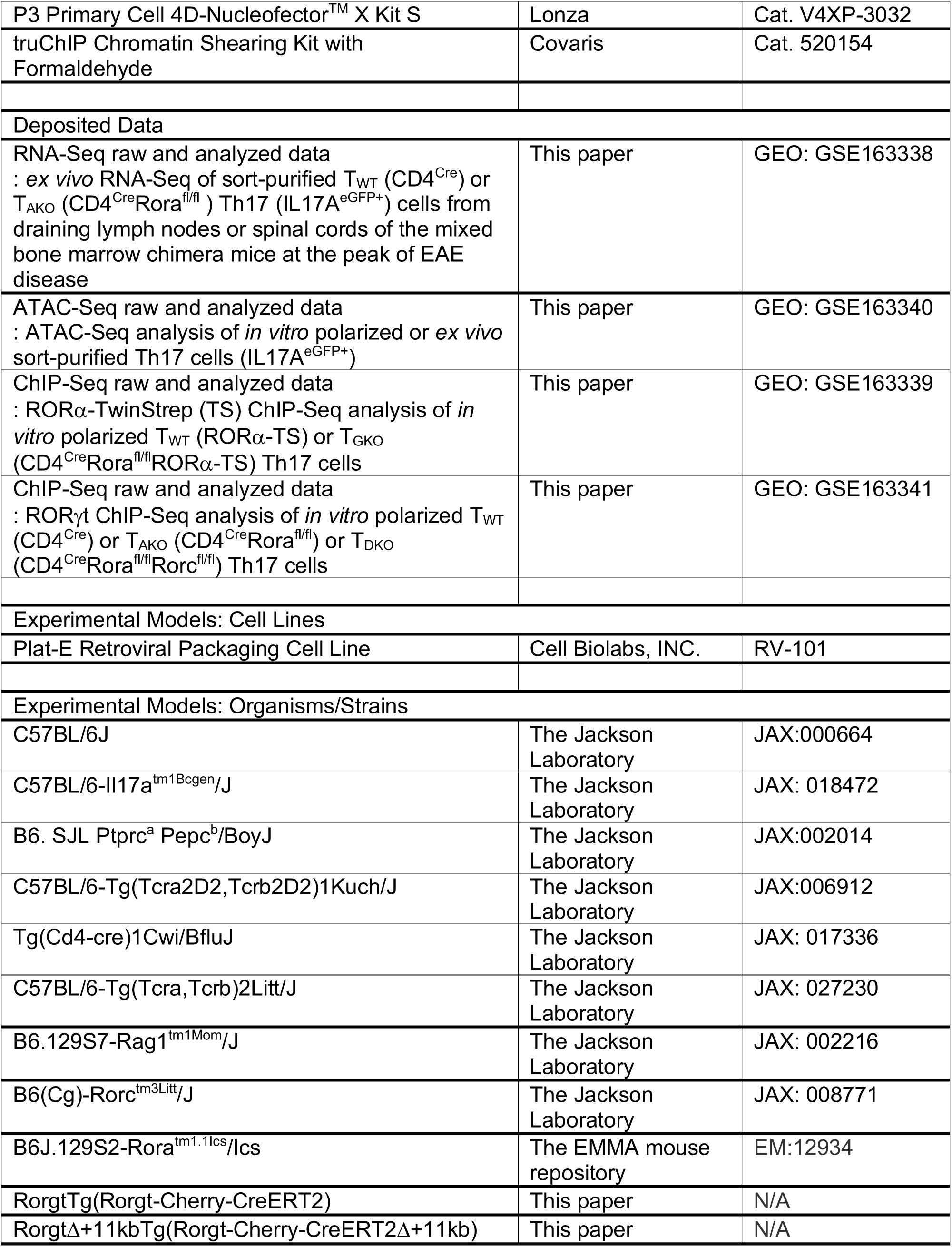
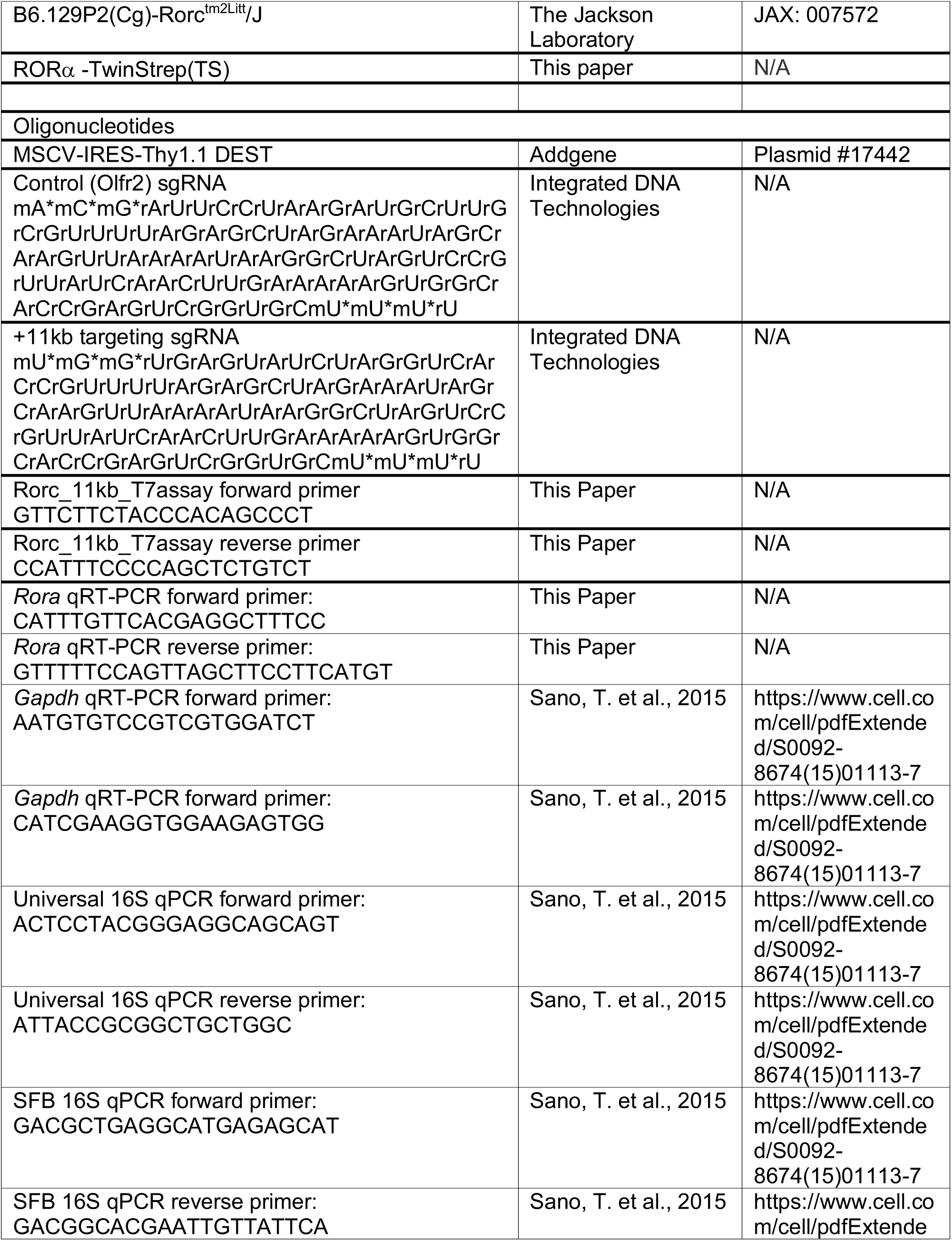
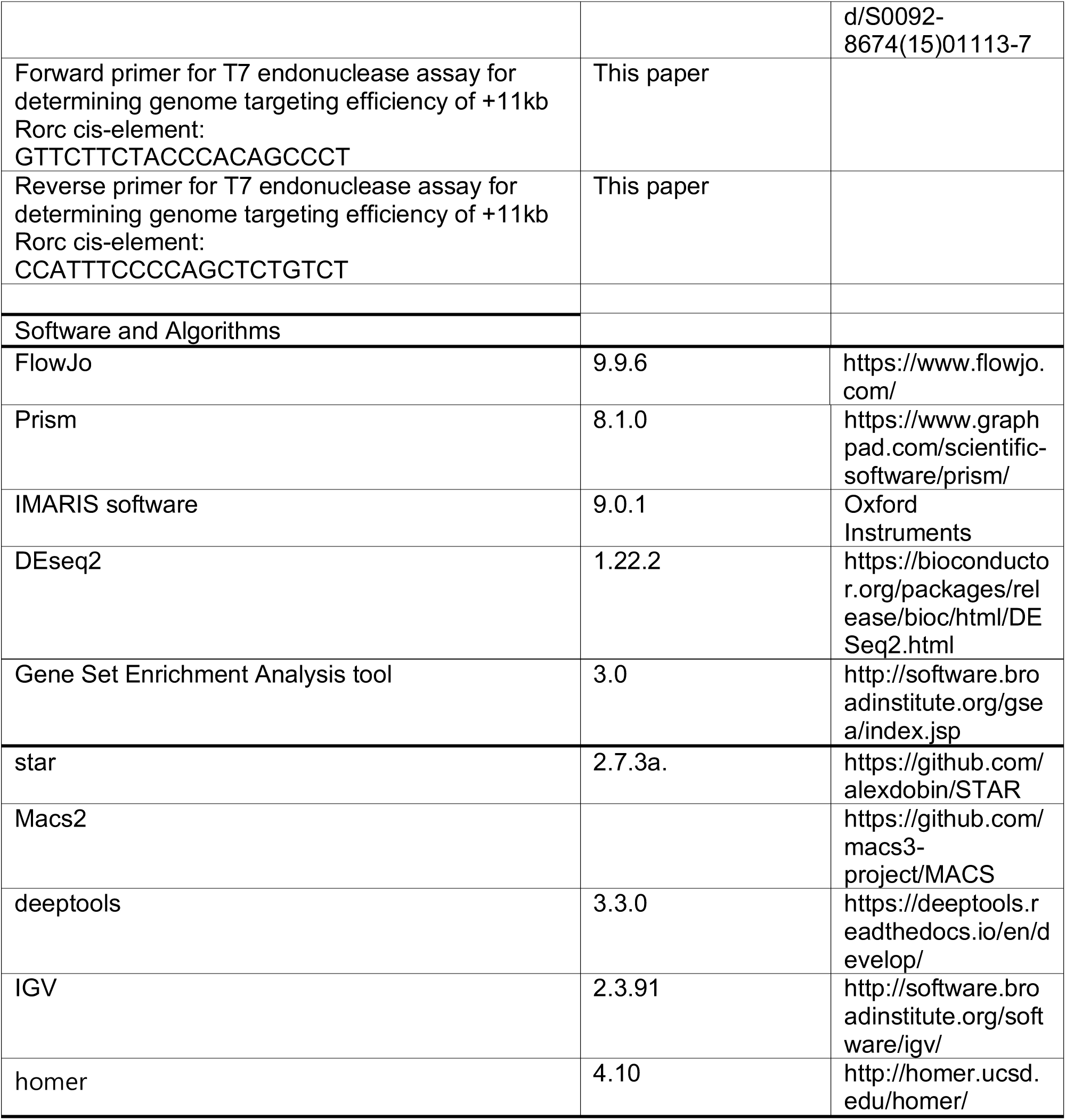

### LEAD CONTACT AND MATERIALS AVAILABILITY

Further information and requests for resources and reagents should be directed to and will be fulfilled by the Lead Contact, Dan R. Littman.

### EXPERIMENTAL MODEL AND SUBJECT DETAILS

#### Mouse Strains

All transgenic animals were bred and maintained in specific-pathogen free (SPF) conditions within the animal facility of the Skirball Institute at NYU School of Medicine. C57BL/6J mice were purchased from The Jackson Laboratory. Frozen sperm of *Rora* “knockout-first” mice (B6J.129S2-Rora^tm1.1Ics^/Ics) mice were obtained from the EMMA mouse repository and rederived onto a C57BL6/J background by NYU School of Medicine’s Rodent Genetic Engineering Core. Wildtype (WT), homozygous Rora floxed (*Rora^fl/fl^*) mice were generated by crossing animals with Tg(Pgk1-flpo)10Sykr mice purchased from The Jackson Laboratories. The flp3 transgene was removed before further breeding to with CD4^Cre^ (Tg(Cd4-cre)1Cwi/BfluJ). *Il17a^eGFP^* reporter (JAX; C57BL/6-Il17a^tm1Bcgen^/J) mice were purchased from The Jackson Laboratories, and bred to the *Rorc* (JAX; B6(Cg)-Rorctm3Litt/J) or *Rora* floxed mutant strains to generate the T_GKO_ (CD4^Cre^Rorgt^fl/fl^) or T_AKO_ (CD4^Cre^Rora^fl/fl^) strains, respectively. T_GKO_ or T_AKO_ strains were further bred to the CD45.1/1 (B6.SJL-Ptprca Pepcb/BoyJ) strain to generated congenically marked lines for co-transfer experiments and mixed bone marrow chimera generation. MOG-specific TCR transgenic (2D2, JAX; C57BL/6-Tg (Tcra2D2,Tcrb2D2)1 Kuch/J) mice were purchased from The Jackson Laboratories, maintained on CD45.1 background, and bred to the T_AKO_ strain. RAG1 knock-out (B6.129S7-Rag1tm1Mom/J) mice were purchased from The Jackson Laboratories, and maintained on CD45.1 background. SFB-specific TCR transgenic (7B8, JAX; C57BL/6-Tg(Tcra,Tcrb)2Litt/J) mice (Yang et al., 2014) were previously described, maintained on an Ly5.1 background, and bred to the T_AKO_ strain. RORA-TS mice were generated using CRISPR-Cas9 technology. Twin-Strep (TS) tag sequence was inserted into the last exon of the *Rora* locus in WT zygotes. Guide RNA and HDR donor template sequences are listed in Table S1. RORA-TS mice were bred with T_GKO_ mice to generate *Rorc* knock-out RORA-TS mice. RorgtTg (Rorgt-Cherry-CreERT2) and RorgtΔ+11kbTg (Rorgt-Cherry-CreERT2Δ+11kb) transgenic reporter mouse lines were generated by random insertion of bacterial artificial chromosomes (BACs) as described below. All in-house developed strains were generated by the Rodent Genetic Engineering Core (RGEC) at NYULMC. Age-(6-12 weeks) and sex-(both males and females) matched littermates stably colonized with Segmented Filamentous Bacteria (SFB) were used for all experiments. To assay SFB colonization, SFB-specific 16S primers were used and universal 16S and/or host genomic DNA were quantified simultaneously to normalize SFB colonization in each sample. All animal procedures were performed in accordance with protocols approved by the Institutional Animal Care and Usage Committee of New York University School of Medicine.

#### Generation of BAC transgenic reporter mice

BAC clone RP24-209K20 was obtained from CHORI (BAC PAC) and BAC DNA was prepared using the BAC100 kit (Clontech). Purified BAC DNA was then electroporated into the recombineering bacterial line SW105. The cassette containing 50bp homology arms surrounding the *Rorc(t)* translational start site ATG was linked to the mCherry-P2A-iCreERT2-FRT-Neo-FRT cassette by cloning into the pL451 vector. The resulting fragment was then excised using restriction digest and gel purified. Homologous recombination was performed by growing the BAC-containing SW105 cells to OD 600 and then heat shocking at 42°C for 15 minutes to induce expression of recombination machinery followed by cooling and washing with H_2_0 to generate electrocompetent cells. These were then electroporated with 0.1μg of purified targeting construct DNA. Correctly recombined bacteria were selected using chloramphenicol (BAC) and Kanamycin. The resultant BAC was purified, screened for integrity of BAC and recombineering junctions by PCR. This BAC was used subsequently to make scarless deletions of putative cis-regulatory elements using GalK positive negative selection according to the Soren Warming protocol #3. The primers, listed in Table S2, were used for generating amplicons for GalK recombineering, and screening for correct insertion and later removal of the GalK cassette.

The primers, listed in Table S3, were used for the recombineering that led to scarless deletion of cis-elements. Correct deletions were confirmed by PCR. The Neo cassette was removed in bacteria via Arabinose inducible Flipase expression and confirmed by PCR. To generate mice, purified BAC DNA was linearized by PI-SceI digestions, dialyzed using Injection buffer (10mM Tris-HCL pH 7.5, 0.1mM EDTA, 100mM NaCl, 30μM spermine, 70μM spermidine) to a concentration of 4ng/μl for microinjection into zygotes.

#### *In vitro* T cell culture and phenotypic analysis

Mouse T cells were purified from lymph nodes and spleens of six to eight week old mice, by sorting live (DAPI^-^), CD4^+^CD25^-^CD62L^+^CD44^low^ naïve T cells using a FACSAria (BD). Detailed antibody information is provided in the Key Resource Table. Cells were cultured in IMDM (Sigma) supplemented with 10% heat-inactivated FBS (Hyclone), 10U/ml penicillin-streptomycin (Invitrogen), 10μg/ml gentamicin (Gibco), 4mM L-glutamine, and 50μM β-mercaptoethanol. For T cell polarization, 1 x 10^5^ cells were seeded in 200μl/well in 96-well plates that were pre-coated with a 1:20 dilution of goat anti-hamster IgG in PBS (STOCK = 1mg/ml, MP Biomedicals Catalog # 55398). Naïve T cells were primed with anti-CD3ε (0.25μg/mL) and anti-CD28 (1μg/mL) for 24 hours prior to polarization. Cells were further cultured for 48h under Th-lineage polarizing conditions; Th0 (Con. : 100U/mL IL-2, 2.5μg/mL anti-IL-4, 2.5μg/mL anti-IFN*γ*), Th17 (0.3 ng/mL TGF-β, 20 ng/mL IL-6, 20 ng/mL IL-23, 2.5μg/mL anti-IL-4, 2.5μg/mL anti-IFN*γ*).

### METHOD DETAILS

#### Flow cytometry

Single cell suspensions were pelleted and resuspended with surface-staining antibodies in HEPES Buffered HBSS containing anti-CD16/anti-CD32. Staining was performed for 20-30min on ice. Surface-stained cells were washed and resuspended in live/dead fixable blue (ThermoFisher) for 5 minutes prior to fixation. PE and APC-conjugated MHC class II (I-A^b^) MOG_38-49_ tetramers (GWYRSPFSRVVH) were provided by the NIH tetramer core facility. PE and APC-conjugated MHC class II (I-A^b^) LT_166-178_ tetramers (RYYRNLNIAPAED) were produced and kindly provided by Timothy Hand’s laboratory at University of Pittsburgh. Staining of tetramer positive T cells was carried out after magnetic isolation of the cells as described (Moon et al., 2009). All tetramer stains were performed at room temperature for 45–60 minutes. For transcription factor staining, cells were treated using the FoxP3 staining buffer set from eBioscience according to the manufacturer’s protocol. Intracellular stains were prepared in 1X eBioscience permwash buffer containing normal mouse IgG (conc), and normal rat IgG (conc). Staining was performed for 30-60min on ice. For cytokine analysis, cells were initially incubated for 3h in RPMI or IMDM with 10% FBS, phorbol 12-myristate 13-acetate (PMA) (50 ng/ml; Sigma), ionomycin (500 ng/ml;Sigma) and GolgiStop (BD). After surface and live/dead staining, cells were treated using the Cytofix/Cytoperm buffer set from BD Biosciences according to the manufacturer’s protocol. Intracellular stains were prepared in BD permwash in the same manner used for transcription factor staining. For EdU staining, we followed manufacturer’s instruction (EdU Flow Cytometry Kit, baseclick). Absolute numbers of isolated cells from peripheral mouse tissues in all studies were determined by comparing the ratio of cell events to bead events of CountBright™ absolute counting beads. Flow cytometric analysis was performed on an LSR II (BD Biosciences) or an Aria II (BD Biosciences) and analyzed using FlowJo software (Tree Star).

#### Induction of EAE by MOG-immunization

For induction of active experimental autoimmune encephalomyelitis (EAE), mice were immunized subcutaneously on day 0 with 100μg of MOG_35-55_ peptide, emulsified in CFA (Complete Freund’s Adjuvant supplemented with 2mg/mL Mycobacterium tuberculosis H37Ra), and injected i.p. on days 0 and 2 with 200 ng pertussis toxin (Calbiochem). For 2D2 transfer EAE experiments, after retrovirus transduction and/or CAS9/RNP electroporation (described below), CD45.1/2 T_WT_ and CD45.2/2 T_AKO_ 2D2 cells were differentiated to ROR*γ*t^+^ effector Th17 cells under the Th17 polarizing condition in vitro for 4 days, then were mixed 1:1 and injected intravenously into recipient mice at total 2 × 10^5^ ROR*γ*t^+^ 2D2 cells per recipient (CD4^Cre^/CD45.1/1). The recipient mice were subsequently immunized for inducing EAE. The EAE scoring system was as follows: 0-no disease, 1-Partially limp tail; 2-Paralyzed tail; 3-Hind limb paresis, uncoordinated movement; 4-One hind limb paralyzed; 5-Both hind limbs paralyzed; 6- Hind limbs paralyzed, weakness in forelimbs; 7- Hind limbs paralyzed, one forelimb paralyzed; 8- Hind limbs paralyzed, both forelimbs paralyzed; 9- Moribund; 10- Death. For isolating mononuclear cells from spinal cords during EAE, spinal cords were mechanically disrupted and dissociated in RPMI containing collagenase (1 mg/ml collagenaseD; Roche), DNase I (100 μg/ml; Sigma) and 10% FBS at 37 °C for 30 min. Leukocytes were collected at the interface of a 40%/80% Percoll gradient (GE Healthcare).

#### Retroviral reconstitution of *Rora* or the RORα -target genes into T_AKO_ 2D2 cells

To generate the ectopic expression retrovirus vector, mouse *Rora*, *Rorc(t)* and *Bhlhe40* were subcloned into the retroviral vector, MSCV-IRES-Thy1.1 (MiT). MiT-Rora, MiT-Rorc(t), MiT- Bhlhe40, and MiT (“empty” vector) plasmids were transfected into PLAT-E retroviral packaging cell line (Cell Bioloab, INC.) using TransIT®-293 transfection reagent (Mirus). Supernatants were collected at 48 h after transfection. Naive T_WT_ or T_AKO_ 2D2 cells were isolated and activated by plate-bound anti-CD3 and anti-CD28. 24 hours after activation, cells were spin-infected by retroviruses MiT-Rora, MiT-Bhlhe40 or control empty vector (MiT-Empty) as described previously (Skon et al., 2013), then were further cultured for 96hrs under Th17- lineage polarizing condition; 20 ng/mL IL-6, 20 ng/mL IL-23, 2.5μg/mL anti-IL-4, 2.5μg/mL anti-IFN*γ*. Prior to adoptive transfer into recipients, Thy1.1+ transduced cells were labeled and enriched with EasySep™ Mouse CD90.1 Positive Selection Kit (STEMCELL).

#### CRISPR mutation of RORE in the +11kb *cis*-element of *Rorc* in 2D2 T cells

To mutate RORE in the +11kb enhancer element of *Rorc*, we delivered CRISPR-Cas9 ribonucleoprotein (RNP) complexes, containing Alt-R CRISPR-Cas9 guide RNAs (the RORE targeting or control sgRNA sequences are listed in the table of STAR Methods) and Cas9 nuclease, into 2D2 cells using electroporation with the Amaxa Nucleofector system (Lonza); 20 μM (1:1.2, Cas9:sgRNA) Alt-R (Integrated DNA Technologies, Inc) Cas9 RNP complex, and 20 μM Alt-R Cas9 Electroporation Enhancer (Integrated DNA Technologies, Inc) as described previously (Vakulskas et al., 2018). sgRNAs were designed using the Crispr guide design software (Integrated DNA Technologies, Inc). FACS-sorted naïve (CD4^+^CD8^-^CD25^-^ CD62L^+^CD44^low^) 2D2 T cells were primed for 18 hrs in T cell medium (RPMI supplemented with 10% FCS, 2mM b-mercaptoethanol, 2mM glutamine), along with anti-CD3 (BioXcell, clone 145- 2C11, 0.25 mg/ml) and anti-CD28 (BioXcell, clone 37.5.1, 1 mg/ml) antibodies on tissue culture plates, coated with polyclonal goat anti-hamster IgG (MP Biomedicals). RNPs were formed by the addition of purified Cas9 protein to sgRNAs in 1 × PBS. Complexes were allowed to form for 30 min at 37°C before electroporation. RNP complexes (5 μL) and 1×10^6^ 2D2 cells (20 μL) were mixed and electroporated according to the manufacturer’s specifications using protocol DN-100 (P3 Primary Cell 4D-Nucleofector^TM^). After 4hrs of recovery in pre-warmed T cell culture medium (Mouse T Cell Nucleofector^TM^ Medium), the electroporated 2D2 cells were polarized into Th17 cells for 96hrs under Th17-lineage polarizing condition; 20 ng/mL IL-6, 20 ng/mL IL- 23, 2.5μg/mL anti-IL-4, 2.5μg/mL anti-IFN*γ*. For *Rora* reconstitution experiment described in Figure S6C, MiT-Rora, MiT-Rorc(t) and MiT (empty) retrovirus were transduced after 24hrs of the electroporation. Prior to adoptive transfer into recipients, Thy1.1^+^ transduced cells were labeled and enriched with EasySep™ Mouse CD90.1 Positive Selection Kit (STEMCELL). The genome targeting efficiency was determined by T7 endonuclease assay (NEB) followed by manufacturer’s protocol (Figure S6A). In parallel, RORE locus of the +11kb cis-element of *Rorc(t)* locus was PCR amplified and cloned into pCR™2.1 vector (ThermoFisher), and mutations in the RORE locus was confirmed by sanger sequencing of the clones (Figure S6B).

#### Generation of bone marrow (BM) chimeric reconstituted mice

Bone marrow (BM) mononuclear cells were isolated from donor mice by flushing the long bones. To generate T_WT_/T_GKO_ chimeric reconstituted mice, CD45.1/2 T_WT_ (*CD4^Cre^Rorc*^+/+^) and CD45.2/2 T_GKO_ (*CD4^Cre^Rorc^fl/fl^*) mice were used as donors. To generate T_WT_/T_AKO_ chimeric reconstituted mice, CD45.1/2 T_WT_ (*CD4^Cre^Rora*^+/+^) and CD45.2/2 T_AKO_ (*CD4^Cre^Rora^fl/fl^*) mice were used as donors. Red blood cells were lysed with ACK Lysing Buffer, and lymphocytes were labeled with Thy1.2 magnetic microbeads and depleted with a Miltenyi LD column. The remaining cells were resuspended in PBS for injection in at 1:4 (T_WT_:T_GKO_) or 1:1 ratio (T_WT_: T_AKO_) to achieve 1:1 chimerism of peripheral T cell populations. Total 5×10^6^ mixed BM cells were injected intravenously into 6 week old RAG1 knock-out recipient mice that were irradiated 4h before reconstitution using 1000 rads/mouse (2×500rads, at an interval of 3h, at X-RAD 320 X-Ray Irradiator). Peripheral blood samples were collected and analyzed by FACS 7 weeks later to check for reconstitution.

#### Oral vaccination

Double mutant E. coli heat labile toxin (R192G/L211A) (dmLT), was produced from *E. coli* clones expressing recombinant protein as previously described (Norton et al., 2011). Mice were immunized twice, 7 days apart by oral gavage, and vaccine responses were assayed 2 weeks after primary gavage as described before (Hall et al., 2008).

#### Isolation of lamina propria lymphocytes

The intestine (small and/or large) was removed immediately after euthanasia, carefully stripped of mesenteric fat and Peyer’s patches/cecal patch, sliced longitudinally and vigorously washed in cold HEPES buffered (25mM), divalent cation-free HBSS to remove all fecal traces. The tissue was cut into 1-inch fragments and placed in a 50ml conical containing 10ml of HEPES buffered (25mM), divalent cation-free HBSS and 1 mM of fresh DTT. The conical was placed in a bacterial shaker set to 37 °C and 200rpm for 10 minutes. After 45 seconds of vigorously shaking the conical by hand, the tissue was moved to a fresh conical containing 10ml of HEPES buffered (25mM), divalent cation-free HBSS and 5 mM of EDTA. The conical was placed in a bacterial shaker set to 37 °C and 200rpm for 10 minutes. After 45 seconds of vigorously shaking the conical by hand, the EDTA wash was repeated once more in order to completely remove epithelial cells. The tissue was minced and digested in 5-7ml of 10% FBS-supplemented RPMI containing collagenase (1 mg/ml collagenaseD; Roche), DNase I (100 μg/ml; Sigma), dispase (0.05 U/ml; Worthington) and subjected to constant shaking at 155rpm, 37 °C for 35 min (small intestine) or 55 min (large intestine). Digested tissue was vigorously shaken by hand for 2 min before adding 2 volumes of media and subsequently passed through a 70 μm cell strainer. The tissue was spun down and resuspended in 40% buffered percoll solution, which was then aliquoted into a 15ml conical. An equal volume of 80% buffered percoll solution was underlaid to create a sharp interface. The tube was spun at 2200rpm for 22 minutes at 22 °C to enrich for live mononuclear cells. Lamina propria (LP) lymphocytes were collected from the interface and washed once prior to staining.

#### SFB-specific T cell proliferation assay

Sorted naive 7B8 or 2D2 CD45.1/1 CD4 T cells were stained with CellTrace™ Violet Cell Proliferation Kit (Life Technology) followed by manufacturer’s protocol. Labeled cells were administered into SFB-colonized congenic CD45.2/2 recipient mouse by i.v. injection. MLNs of the SFB-colonized mice were collected at 96h post transfer for cell division analysis.

#### RNA isolation and library preparation for RNA sequencing

Total RNAs from in vitro polarized T cells or sorted cell populations were extracted using TRIzol (Invitrogen) followed by DNase I (Qiagen) treatment and cleanup with RNeasy MinElute kit (Qiagen) following manufacturer protocols. RNA-Seq libraries for *ex vivo* isolated IL17^eGFP+^ T_WT_ or T_AKO_ Th17 lineages from DLN or spinal cords of immunized BM chimeras at peak of EAE were prepared with the SMART-Seq® v4 PLUS Kit (Takara, R400752). The sequencing was performed using the Illumina NovaSeq or NextSeq. RNA-seq libraries were prepared and sequenced by the Genome Technology Core at New York University School of Medicine.

#### Library preparation for ATAC sequencing

Samples were prepared as previously described (Buenrostro et al., 2013). Briefly, 50,000 sort-purified Th17 cells were pelleted in a fixed rotor centrifuge at 500xg for 5 minutes, washed once with 50 μL of cold 1x PBS buffer. Spun down again at 500xg for 5 min. Cells were gently pipetted to resuspend the cell pellet in 50 μL of cold lysis buffer (10 mM Tris-HCl, pH7.4, 10 mM NaCl, 3 mM MgCl2, 0.1% IGEPAL CA-630) for 10 minutes. Cells were then spun down immediately at 500xg for 10 min and 4 degrees after which the supernatant was discarded and proceeded immediately to the Tn5 transposition reaction. Gently pipette to resuspend nuclei in the transposition reaction mix. Incubate the transposition reaction at 37 degrees for 30 min. Immediately following transposition, purify using a Qiagen MinElute Kit. Elute transposed DNA in 10 μL Elution Buffer (10mM Tris buffer, pH 8). Purified DNA can be stored at −20 degrees C. The transposed nuclei were then amplified using NEBNext High-fidelity 2X PCR master mix for 5 cycles. In order to reduce GC and size bias in PCR, the PCR reaction is monitored using qPCR to stop amplification prior to saturation using a qPCR side reaction. The additional number of cycles needed for the remaining 45 μL PCR reaction is determined as following: (1) Plot linear Rn vs. Cycle (2) Set 5000 RF threshold (3) Calculate the # of cycle that is corresponded to ¼ of maximum fluorescent intensity. Purify amplified library using Qiagen PCR Cleanup Kit. Elute the purified library in 20 μL Elution Buffer (10mM Tris Buffer, pH 8). Be sure to dry the column before adding elution buffer. The purified libraries were then run on a high sensitivity Tapestation to determine if proper tagmentation was achieved (band pattern, not too much large untagmented DNA or small overtagmented DNA at the top or bottom of gel. Paired-end 50bp sequences were generated from samples on an Illumina HiSeq2500.

#### Library preparation for Chromatin Immunoprecipitation (ChIP-Seq)

RORα-TS and RORγt ChIP-Seq was performed as described (Ciofani et al., 2012) with the following modifications. For each ChIP, 20-80 million cells were cross-linked with paraformaldehyde; chromatin was isolated using truChIP Chromatin Shearing Kit (Covaris) and fragmented with a S220 Focused-ultrasonicator (Covaris). Twin-strep (TS) tagged RORα protein was precipitated using Strep-TactinXT according to the manufacturer’s protocol (IBA Lifesciences). Following immunoprecipitation, the protein-DNA crosslinks were reversed and DNA was purified. DNA from control samples was prepared similarly but without immunoprecipitation. Sequencing libraries were made from the resulting DNA fragments for both ChIP and controls using DNA SMART™ ChIP-Seq Kit (Takara) for RORα-TS ChIP-Seq and KAPA HyperPlus Kit (Roche) for RORγt ChIP-Seq. The ChIP-Seq libraries were sequenced with paired-end 50 bp reads on an Illumina HiSeq 4000.

### QUANTIFICATION AND STATISTICAL ANALYSIS

#### Transcriptome analysis

##### RNA-Seq methods

Bulk RNA-Seq fastq files were aligned to the mm10 reference genome using star v 2.7.3a. Bam files were converted to bigwig files via deeptools v 3.3.0 bamCoverage for visualization. DEseq2 was used for differential gene analysis.

##### ChIP-Seq methods

ChIP-Seq fastq files were aligned to the mm10 reference genome using star v 2.7.3a. Bam files were converted to bigwig files via deeptools v 3.3.0 bamCoverage and normalized by RPGC to compare peak heights across samples. Deeptools computeMatrix and plotHeatmap were used to make heatmaps. Macs2 was used to call peaks using a significance cutoff of 0.01 for the previously published RORγt ChIP-Seq dataset (Ciofani et al., 2012) (Figure S4E, left panel), 0.5 for the RORα-Twin Strep ChIP-Seq datasets (Figure S4E, middle and right panels), and 0.05 for the RORγt ChIP-Seq datasets (Figure S4I). During peak calling the treatment file was used with its associated control file. The homer annotatePeaks.pl script was used to annotate peaks within 10kb of a gene.

##### ATAC-Seq methods

Bowtie2 was used to align the reads to the mm10 genome using parameters - very-sensitive. Picard tools was used to mark and remove duplicates. Deeptools bamCoverage was used to generate a bigwig file normalized using RPGC.

#### Statistical analysis

Differences between groups were calculated using the unpaired two-sided Welch’s t-test or the two-stage step-up method of Benjamini, Krieger and Yekutieliun. For EAE disease induction, log-rank test using the Mantal-Cox method was performed. For RNA-seq analysis, differentially expressed genes were calculated in DESeq2 using the Wald test with Benjamini–Hochberg correction to determine the FDR. Genes were considered differentially expressed with FDR < 0.01 and log2 fold change > 1.2. Data was processed with GraphPad Prism, Version 8 (GraphPad Software). We treated less than 0.05 of p value as significant differences. ∗p < 0.05, ∗∗p < 0.01, ∗∗∗p < 0.001, and ∗∗∗∗p < 0.0001. Details regarding number of replicates and the definition of center/error bars can be found in figure legends.

## DATA AND CODE AVAILABILITY

The RNA-Seq, ATAC-Seq, ChIP-Seq datasets generated during this study are available at Gene Expression Omnibus (GSE163338, GSE163340, GSE163339, GSE163341).

## Supplemental Tables

**Table S1.**
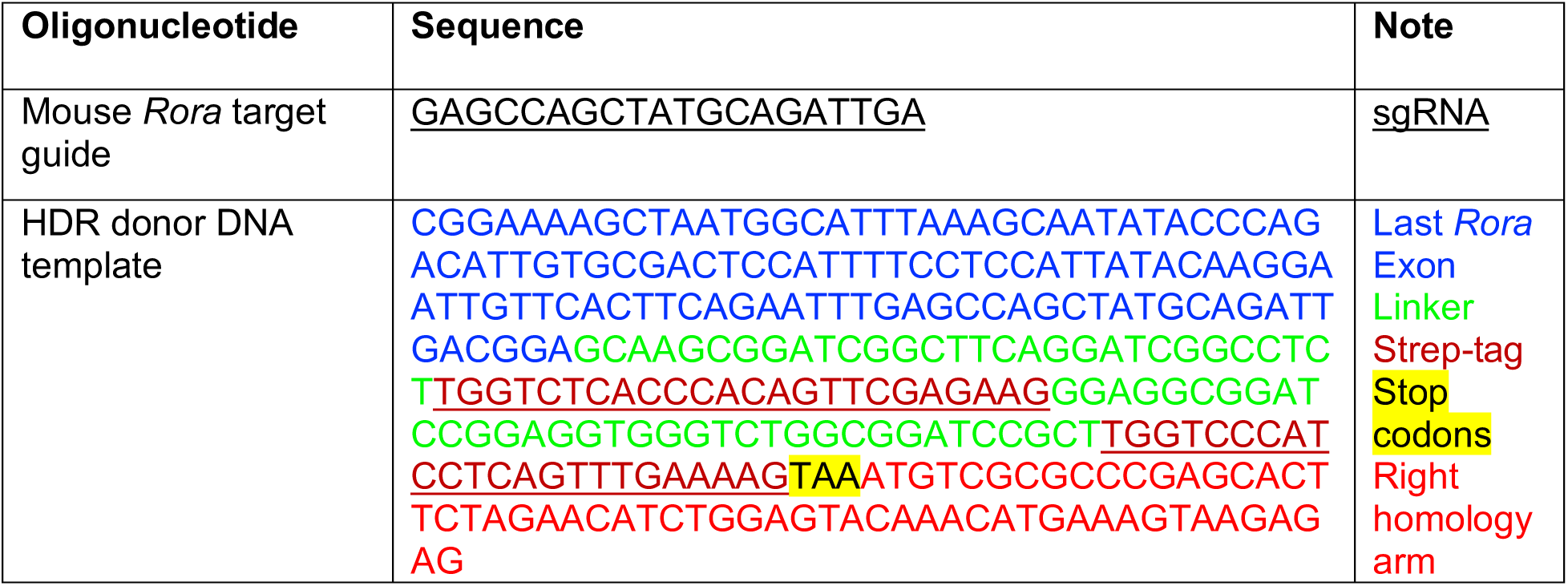
sgRNA and HDR donor sequence, Related to STAR methods.

**Table S2.**
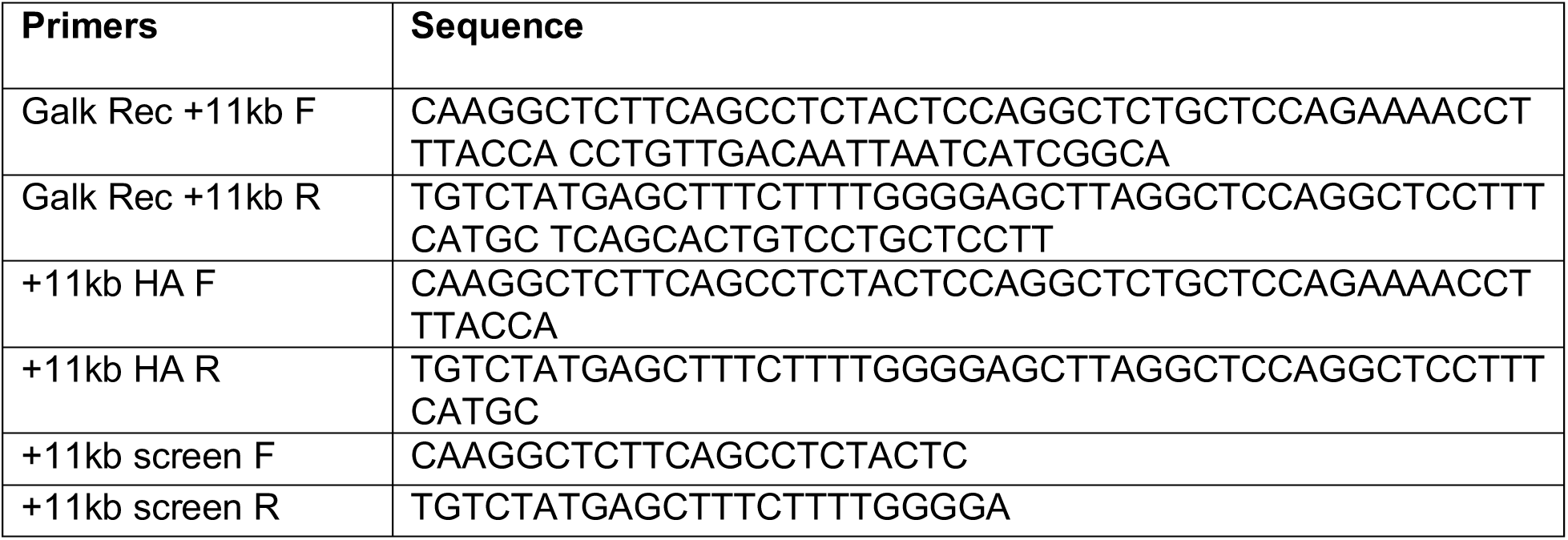
Primers for GalK recombineering and screening.

**Table S3.**
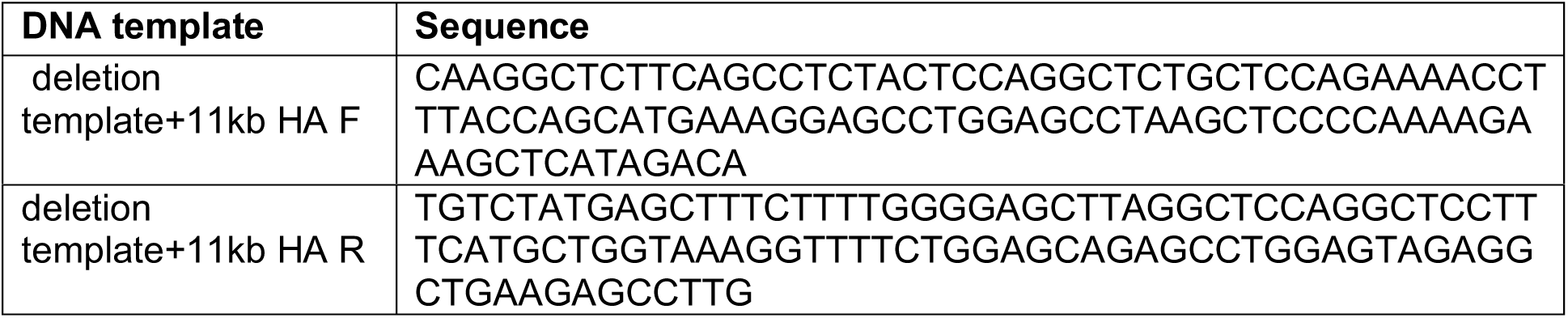
DNA template for *Rorc* deletions.

## Supplemental figure legends

**Figure S1:**
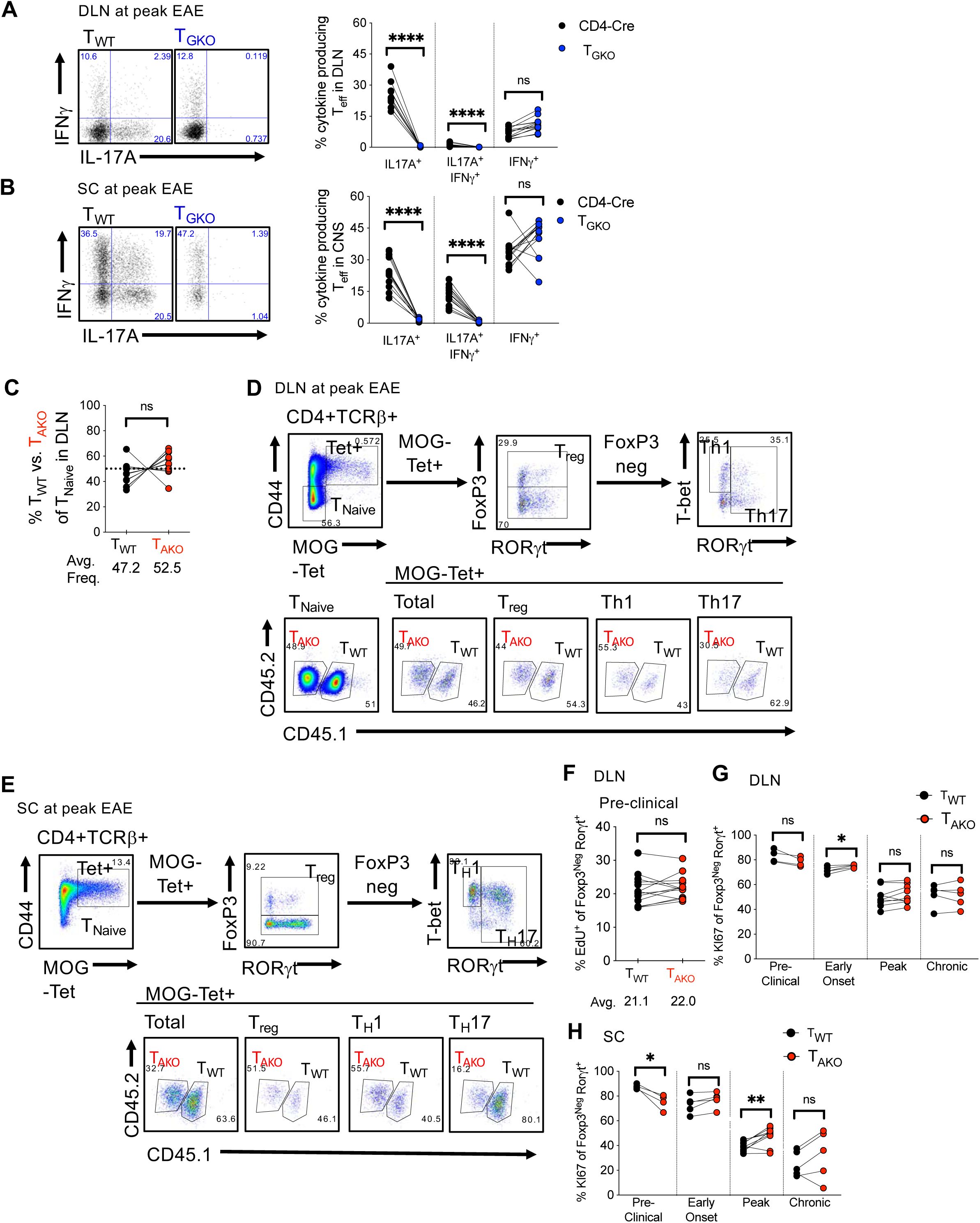
Role of RORγt and RORα in Th17 differentiation and accumulation during autoimmune encephalomyelitis, Related to Figure 1. **(A and B)** IL-17A and IFNγ production of CD44^hi^ effector T cells upon their ex vivo PMA/Iono restimulation. Cells from DLN **(A)** and SC **(B)** of T_WT_/T_GKO_ BM chimera at peak of EAE. Data combined three experiments with 13 BM chimera mice. **(C)** Mean percent donor-derived CD44^lo^ CD4^+^ naïve T cell chimerism at peak of EAE, as determined by flow cytometric analysis of DLN. Data combined three experiments with 12 T_WT_/T_AKO_ BM chimera mice. **(D and E)** Gating strategies to identify all Th populations amongst MOG-tetramer^+^ T_WT_ and T_AKO_ donor-derived CD4^+^ T cells in the DLN (D) and SC (E) of T_WT_/T_AKO_ BM chimera mice at peak of EAE. **(F)** Percent of EdU-incorporating Th17 (RORγt^+^FoxP3^neg^) cells from DLN of T_WT_/T_AKO_ BM chimera mice at pre-clinical stage of EAE. Data combined with 13 T_WT_/T_AKO_ BM chimera mice. **(G and H)** Percent of Ki-67^+^ Th17 (RORγt^+^/FoxP3^neg^) cells from DLN (G) and SC (H) of T_WT_/T_AKO_ BM chimera mice at indicated stages of EAE. Data combined two experiments for the pre-clinical (n=4), early onset (n=5), acute (n=9), and chronic stages (n=5) of disease, respectively. Statistics were calculated using the paired sample T test. ns = not significant, *p < 0.05, **p < 0.01, ****p < 0.0001.

**Figure S2.**
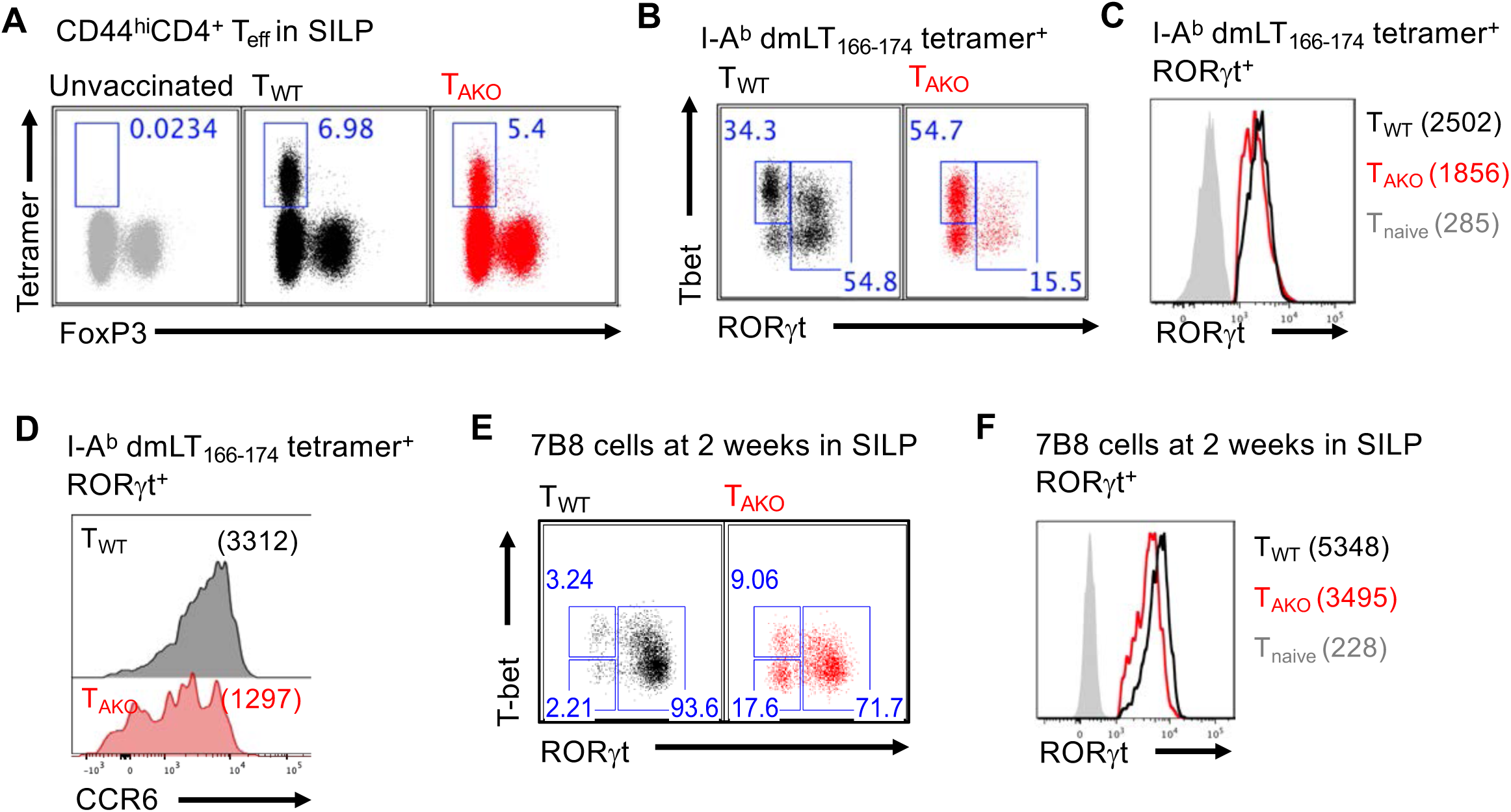
RORα deficiency impairs Th17 cell accumulation in SILP. Related to Figure 2. **(A)** Small intestinal lamina propria CD4^+^CD44^+^ T cells were stained for I-A^b^ dmLT_166-174_ tetramer binding and Foxp3 expression to compare the dmLT-specific CD4^+^ T cell effector responses between T_WT_ and T_AKO_ mice. **(B)** Gated dmLT tetramer^+^ T cells from representative T_WT_ (black dot plot) and T_AKO_ (red dot plot) SILP were analyzed for expression of T-bet and RORγt. **(C and D)** Histograms depicting expression of RORγt (C) and CCR6 (D) in T_WT_ and T_AKO_ dmLT tetramer^+^ RORγt^+^ Th17 cells. Geometric mean fluorescence intensities (gMFI) are included in parentheses. **(E)** Representative flow cytometric analysis of SILP-accumulated T_WT_ (black dot plot) and T_AKO_ (red dot plot) 7B8tg cells at 2 weeks post adoptive transfer. **(F)** Histogram of RORγt expression in T_WT_ and T_AKO_ RORγt^+^ 7B8 Th17 cells. Geometric mean fluorescence intensities (gMFI) are included in parentheses.

**Figure S3.**
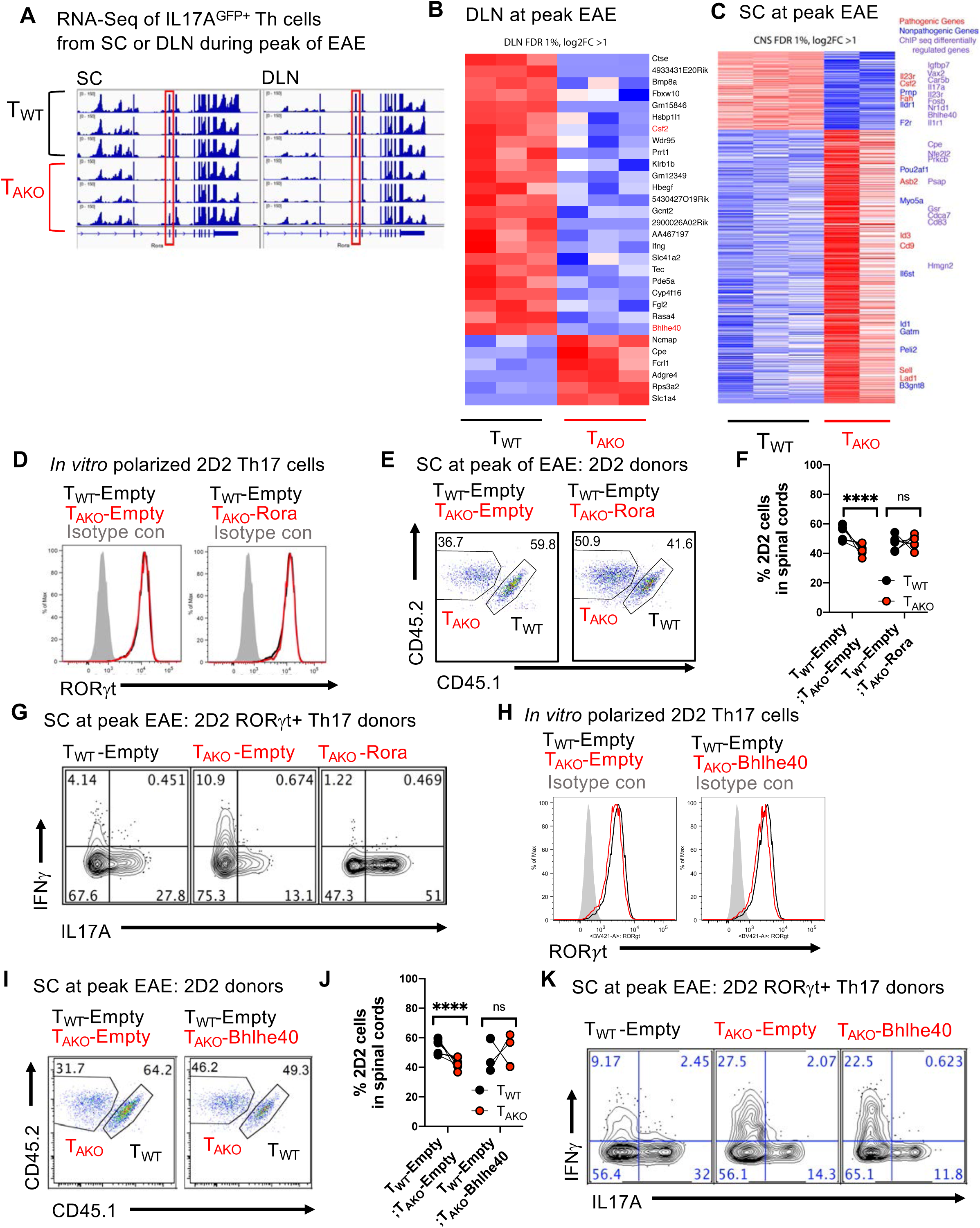
Target genes of RORα, and rescue of EAE phenotype with T_AKO_ cells expressing ectopic RORα or its target gene product Bhlhe40. Related to Figure 3. **(A-C)** RNA-Seq analysis to identify target genes of ROR*α*. RNA preparation from sorted *Il17a^eGFP^*^+^ mice is described in Methods. One SC T_AKO_ Th17 sample contained reads in the deleted region of the *Rora* locus and thus was excluded from analysis; all other T_AKO_ samples were devoid of reads in this region (A). (A) RNA-Seq tracks within *Rora* locus indicating efficient inducible deletion of *Rora* (Exon3) of *Il17a^eGFP^*^+^ T_WT_ and T_AKO_ Th17 cells from DLN and SC of mixed BM chimera mice at peak of EAE. (B and C) Clustered heatmap of differentially expressed genes between *Il17a^eGFP^*^+^ Th17 T_WT_ and T_AKO_ cells from DLN (B) and SC (C) of mixed BM chimera mice at peak of EAE. Color scale is based on z-scores for each gene. Genes listed on the righthand margin are color coded. Blue = non-pathogenic Th17 signature. Red = pathogenic Th17 signature. Purple = Genes associated with RORα ChIP-Seq peaks. **(D-K)** Reconstitution of 2D2 T_AKO_ cells with RORα or Bhlhe40 and phenotypic analysis in spinal cords during EAE. (D) Stacked histogram illustrating representative RORγt expression of *in vitro* polarized 2D2tg T_WT_-Empty and RORα-deficient (T_AKO_-Empty) or -reconstituted (T_AKO_-Rora) Th17 cells. (E and F) Representative FACS plots (E) and frequency (F) of co-transferred T_WT_ and T_AKO_ donor-derived 2D2tg cells, retrovirally reconstituted with or without *Rora*, in the SC at peak of EAE. (G) Representative FACS plots displaying IL-17A and IFNγ production of RORγt^+^ Th17 T_AKO_-Empty or T_AKO_-Rora 2D2 cells compared to T_WT_-Empty upon ex vivo PMA/Ionomycin restimulation. (H) Stacked histogram illustrating representative RORγt expression of *in vitro* polarized 2D2tg T_WT_-Empty and Rora-deficient (T_AKO_-Empty) or Bhlhe40-overexpressing (T_AKO_- Bhlhe40) Th17 cells. (I and J) Representative FACS plots (I) and frequencies (J) of co-transferred T_WT_ and T_AKO_ donor-derived 2D2tg cells, retrovirally transduced with or without *Bhlhe40*, in the SC at peak of EAE. (K) Representative FACS plots displaying IL-17A and IFNγ production of RORγt^+^ Th17 T_AKO_-Empty or T_AKO_-Bhlhe40 2D2 cells compared to T_WT_-Empty upon ex vivo PMA/Ionomycin re-stimulation. Data combined from two experiments. Statistics were calculated using the paired sample T test. ns = not significant, *p < 0.05, **p < 0.01, ***p < 0.001, ****p < 0.0001.

**Figure S4.**
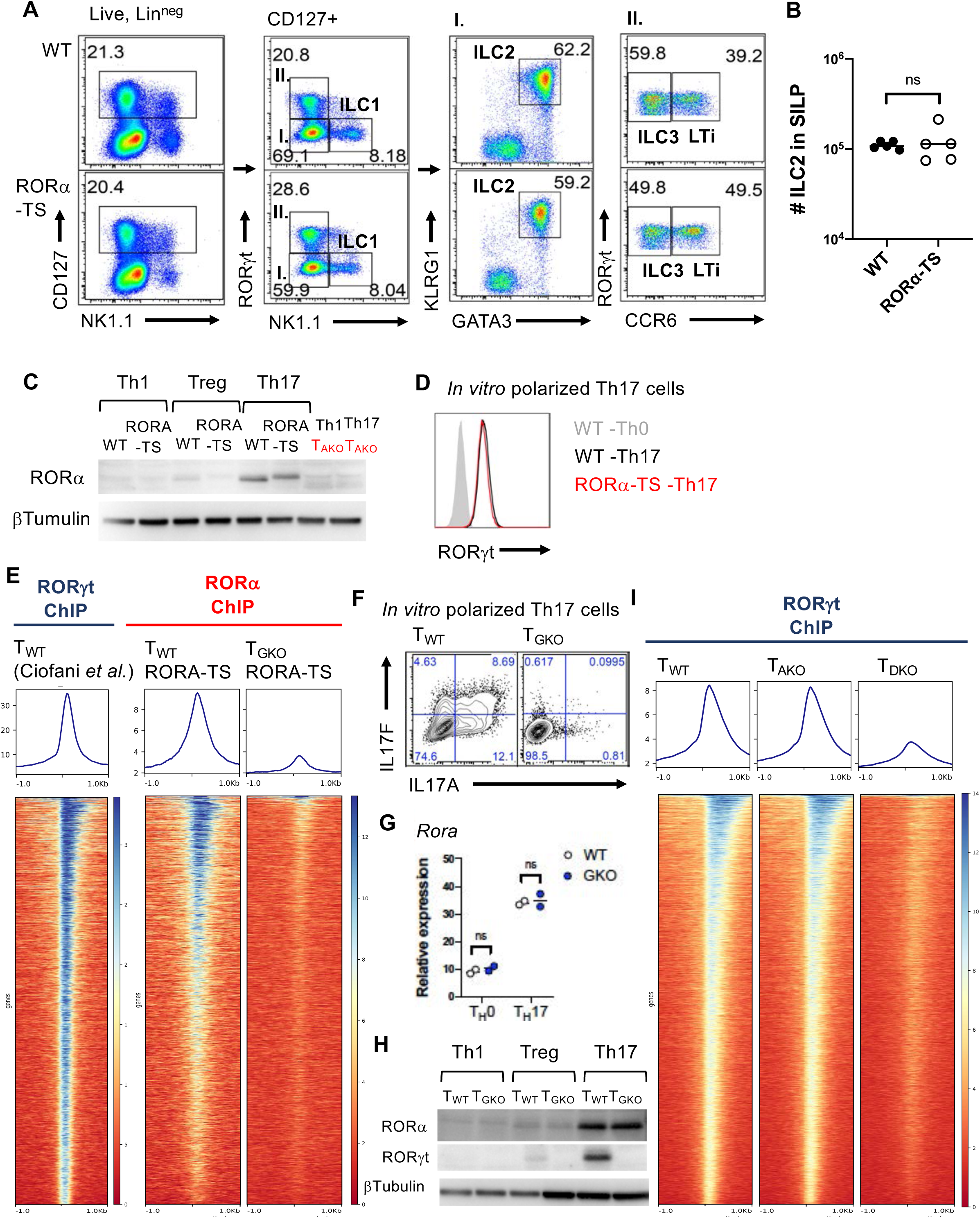
Shared genomic binding sites of RORα and RORγt in Th17 cells. Related to Figure 4. **(A)** Gating strategy to identify innate lymphoid cell (ILC) populations in small intestine lamina propria (SILP) of wild type (WT) and RORA-TS mice. Lineage markers (Lin) include CD3, TCRβ, TCRγδ, CD11b, CD19. **(B)** Absolute number of ILC2 (Lin^neg^, CD127^+^, RORγt^neg^, NK1.1^neg^, KLRG1^+^, GATA3^+^) in SILP of WT and RORA-TS mice. **(C)** Western blot data displaying intact RORα expression of *in vitro* polarized RORA-TS Th17 cells. **(D)** Stacked histogram illustrates representative RORγt expression of *in vitro* polarized RORA- TS Th17 cells. **(E)** Heatmaps depicting genome-wide RORγt (left) and RORα-TS (middle and right) ChIP-Seq signals of *in vitro* polarized Th17 cells, centered on the summit of RORγt binding sites called on the basis of our earlier dataset (Ciofani et al., 2012). Middle and right alignments compare RORα occupancy in wild-type and RORγt-deficient T cells. **(F)** Representative FACS plots displaying IL-17A and IL-17F production of *in vitro* polarized T_WT_ or T_GKO_ Th17 cells. **(G)** qPCR result of *Rora* gene expression of in vitro polarized T_WT_ and T_GKO_ cells cultured under Th0 (IL2) or Th17 (IL-6+TGF-β+IL-23) conditions for 48h. **(H)** Immunoblots for RORα and RORγt of in vitro polarized T_WT_ and T_GKO_ cells cultured under Th1 (IL-2+IL-12), Treg (IL-2+TGF-β) or Th17 (IL-6+TGF-β+IL-23) conditions for 48h. β-Tubulin is shown as a loading control. **(I)** Heatmaps representing RORγt ChIP-Seq peaks of *in vitro* polarized T_WT_ (left), T_AKO_ (middle) and RORα/RORγt double knock-out (T_DKO_) (right) Th17 cells.

**Figure S5.**
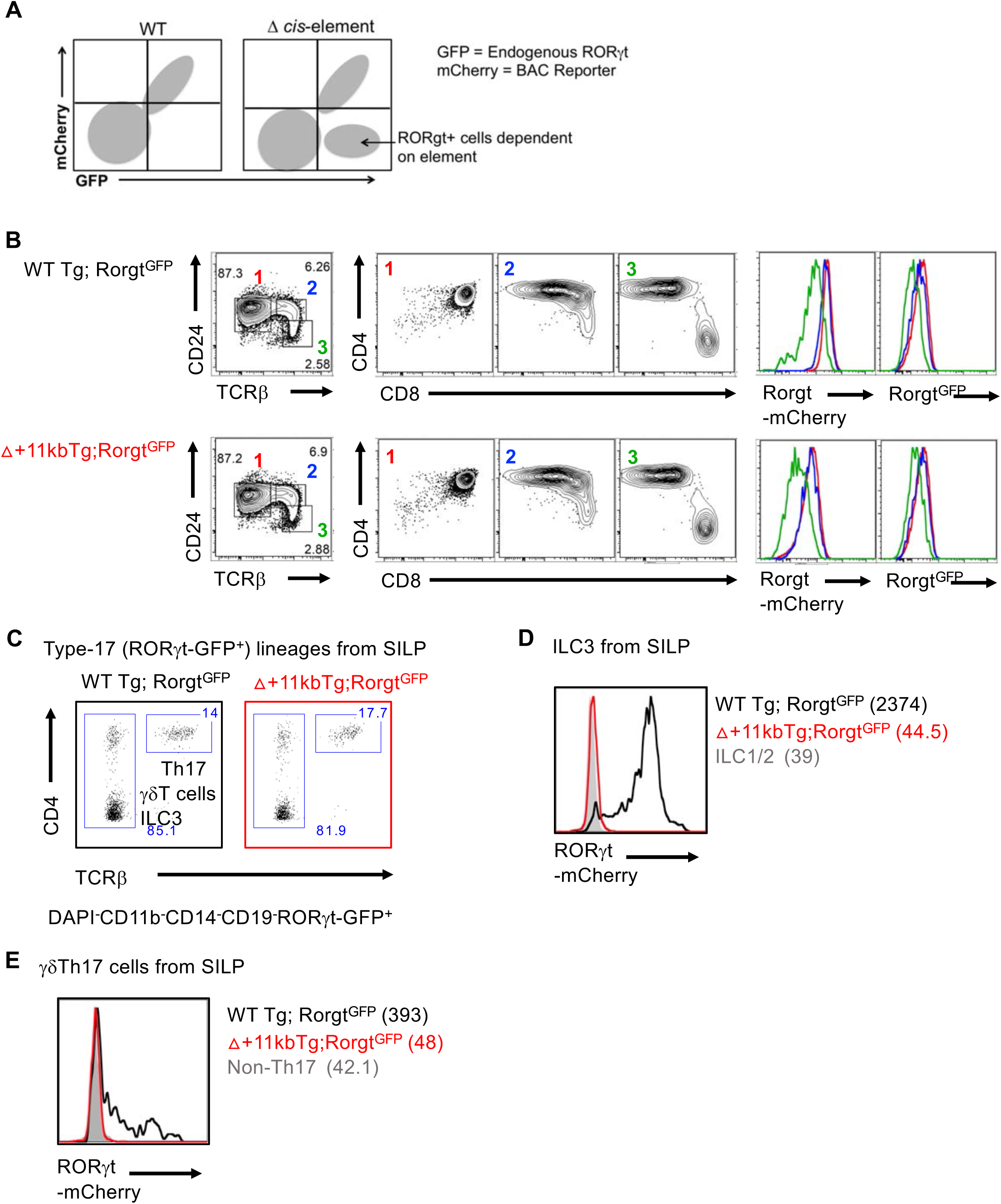
Requirement of *Rorc(t)* +11kb *cis*-element for RORγt expression in Type-17 lymphocytes *in vivo*. Related to Figure 5. **(A)** Schematic depicting expression of endogenous and Tg reporter alleles in *Rorc(t)-mCherry* BAC Tg; *Rorc(t)^GFP^* mice. **(B)** Flow cytometry plots depicting gating strategy to capture thymocyte development from DP (CD4^+^CD8^+^) stage to post-selection stages (left and middle). On the right, mCherry and GFP reporter expression in each color-coded thymocyte subset from indicated Tg mouse line. **(C)** Flow cytometry of indicated populations from the SILP in WT and +11kb enhancer mutant Tg (Δ+11kbTg); *Rorc(t)^GFP^* mice. **(D and E)** mCherry reporter expression of *ex vivo* isolated ILC3 (Lin^neg^RORγt^GFP+^) cells (D) and γδTh17 (γδTCR^+^RORγt^GFP+^) cells (E) from SILP of WT Tg; *Rorc(t)^GFP^* or Δ+11kbTg;*Rorc(t)^GFP^* mice. gMFIs are included in parentheses. Representative data of three experiments.

**Figure S6.**
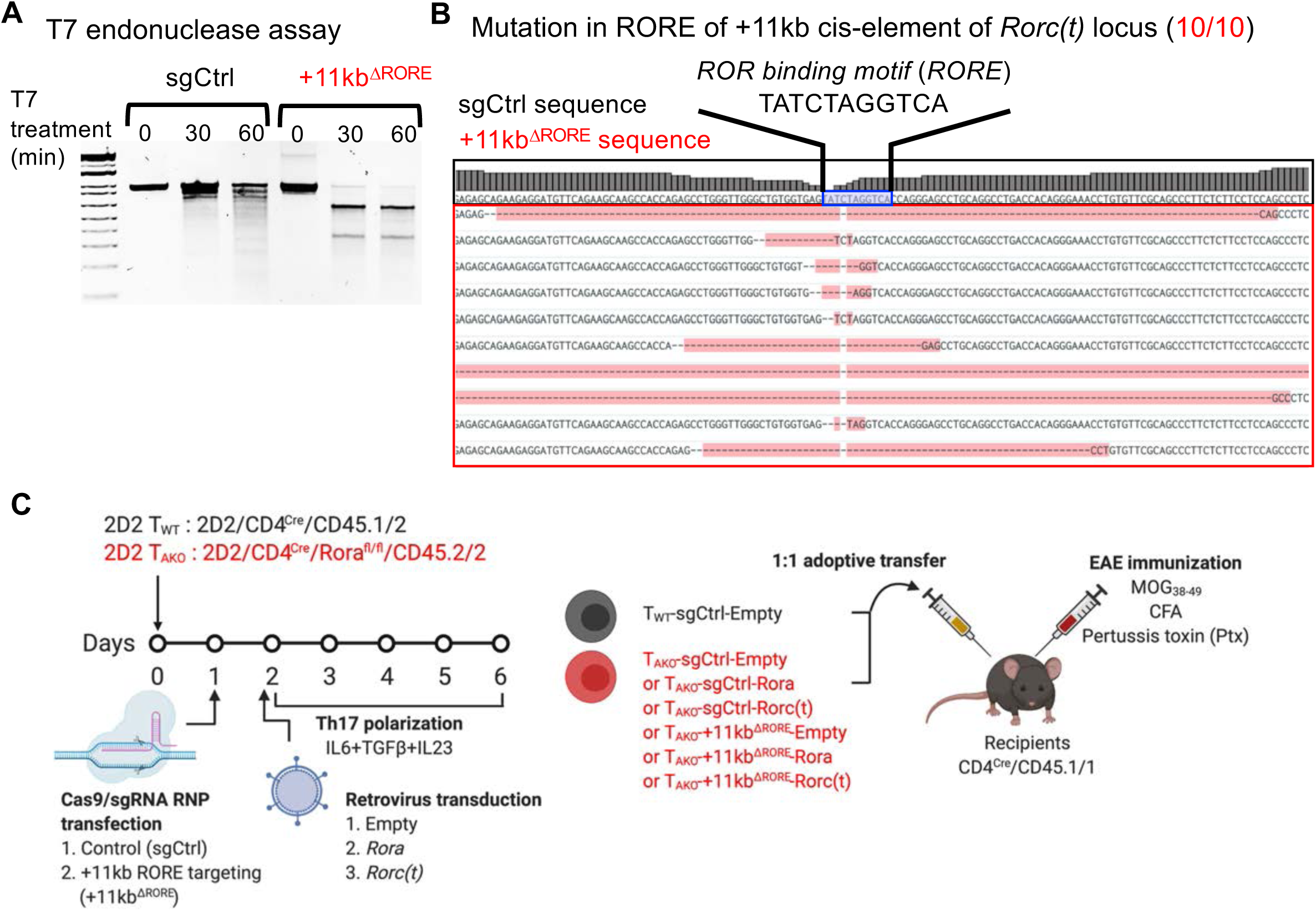
Efficient editing of *Rorc(t)* +11kb *cis*-element by CAS9-RNP method. Related to Figure 6. **(A)** Analysis of CAS9/gRNA RNP-mediated targeting efficiency of +11kb enhancer by T7 endonuclease I assay. **(B)** Sanger sequencing results displaying *Rorc(t)* +11kb enhancer mutations and deletions of T_AKO_ +11kb^ΔRORE^ 2D2tg-Th17 cells. **(C)** Experimental scheme to examine the role of *Rorc(t)* +11kb *cis*-element in maintenance of pathogenic Th17 program during EAE.

